# Is Britain still Great for Pine marten? A Habitat Suitability Assessment

**DOI:** 10.1101/2025.03.19.643030

**Authors:** Ella Lewis, Amy Fitzmaurice, Rachel Gardner, Suzanne Kynaston, Neal Armour-Chelu, Dave J.I. Seaman, Kirsty Swinnerton, Lawrence Ball

## Abstract

**Context:** Species distribution models are used to predict habitat suitability for a species, by quantifying the environmental characteristics that allow a species to occupy a geographical area. The abundance and range of pine marten *(Martes martes)* has declined substantially in Great Britain, with remaining populations restricted to Scotland.

**Objectives:** Here, we perform species distribution modelling using BIOMOD2 platform to determine habitat suitability, and inform the identification of potential reintroduction sites for pine marten in Great Britain.

**Methods:** Using a global range dataset of 4,189 occurrences and seven environmental variables, ensemble species distribution models were used to predict habitat suitability across Europe at 1 km resolution and Great Britain at 100 m resolution.

**Results:** Across the extent of both Europe and Britain, results indicate high suitability in areas with woody vegetation cover in low topographic positions, and notably low in urban areas and extensive areas of arable land. In Britain, high habitat suitability is identified across substantial areas in the South East of England, parts of South West of England, East Yorkshire and Gloucestershire, with pockets of suitable habitat along the West Coast of Britain. The results indicate that elevation and land cover are important drivers of suitability.

**Conclusion:** Habitat suitability modelling at a high resolution of 100m proves effective for informing potential reintroduction sites for pine marten in Britain. We also demonstrate the importance of using occurrence data from pine martens’ global range to predict optimal habitat suitability.

## Introduction

The Pine marten *(Martes martes)* is a mustelid species that typically occupies woodland habitats, distributed across mainland Europe extending to Western Siberia (Buskirk, 1992; Delibes,1983). In Britain, pine marten were once abundant, however their population suffered severe decline during the late 1800s due to habitat loss and hunting practices, particularly predator control for game birds (Langley and Yalden, 1977). The species is classified as critically endangered in England and Wales according to the Red List for British Mammals (Mathews and Harrower, 2020). It is a priority species in the UK Biodiversity Action Plan and has been fully protected since 1988 under Schedule 5 of the Wildlife and Countryside Act 1981. The fragmented populations that have persisted are restricted to north-west Scotland and the north of England and Wales. As an opportunistic omnivore, their diet is comprised of small mammals, particularly voles, small birds, berries and insects (Twining et al., 2019; Jedrzejewski et al., 1993). They play an important role within a woodland ecosystem by providing several functions including seed dispersal and balancing prey populations as a mesopredator (Paine, 1969; Hastings et al., 2007). For example, in Scotland and Ireland, they have been shown to suppress non-native invasive grey squirrel (*Sciurus carolinensis*) populations (Sheehy et al., 2018; Vincent Wildlife Trust, 2020, Reilly and Lawton, 2025). This in turn, allows indirect benefits for vegetation recovery due to reductions in tree damage, and red squirrel (*Sciurus vulgaris*) population recovery by reducing interspecific competition with grey squirrel (Sheehy et al., 2018; Schaumann and Heinken, 2002; Crowley et al., 2018).

Pine martens are considered an indicator species for woodland ecosystems where their presence can denote a biologically diverse ecosystem in good condition (Buskirk, 1992; DEFRA, 2021). As a flagship species, pine marten can act as an effective conservation tool, not only for reintroduction approaches, but also to promote conservation of woodland habitats.

Woodland, defined as land with at least 20% tree canopy cover (Forestry Commission, 2024), accounts for only 13% of land in Britain (Forest Research, 2022), compared to the European average of 37% (Forestry Research 2010), therefore woodland creation and restoration is a national priority (Defra, 2022).

Reintroduction projects can be a key tool for species recovery, and pine marten reintroductions have already occurred across Britain in Gloucestershire, Wales, Devon and Cumbria (MacPherson et al., 2014; Stringer et al., 2015). Reintroductions require extensive research, primarily around social and ecological feasibility pre-release, population genetics, disease risk, dispersal potential, and post-release monitoring (Kirkwood, 2013; Klein and Arts, 2021; Armstrong and Seddon, 2008, Seddon and Armstrong, 2019; IUCN, 2013).

Species distribution models (SDM), sometimes referred to as environmental niche models (ENM) or habitat suitability models (HSM), are a useful tool to identify suitable habitats or environments for a species. SDMs estimate the relationship between observed species occurrences and the environmental conditions at these locations, to characterise the environmental conditions that are suitable for that species. These estimated relationships are then used to identify where suitable environments are distributed in space, in the form of a habitat suitability map (Anderson et al., 2003; Elith and Leathwick, 2009).

A key application for SDMs is to identify sites with high suitability for a species, as an informative tool for conservation planning (Guisan et al., 2006). For example, they can be used to assist in identifying priority areas for conservation management that will have the most impact, based on species richness and suitability. SDMs are also used to predict species’ distributional changes over time and space (Mammola et al., 2021), for instance, to understand how species will respond to future climatic conditions (Raes and Aguirre-Gutiérrez, 2018).

SDMs generally use a single statistical technique to predict habitat suitability for a species. However, the field of habitat suitability mapping has evolved rapidly, giving access to more complex and potentially reliable approaches (Meller et al., 2014). One approach is ensemble species distribution models (eSDMs), which use an assemblage of statistical models to produce suitability predictions. This method has been shown to improve the accuracy of model predictions compared to individual models (Kwon, 2014; Araújo and New, 2007; Hao et al., 2020). Given that there is often significant variation in the predictions across model techniques and there is no single outstanding model algorithm, it is sensible to combine several, well-performing techniques in order to generate accurate predictions (Meller et al., 2014; Li and Wang, 2012). In addition, ensemble forecasting has become more widely used for its ability to combat the limitations of individual algorithms, whilst allowing for comparisons between the performance of different algorithms (Araújo & New, 2007).

Here, we aim to use eSDMs to identify areas in Britain with sufficient suitable habitat to support viable breeding populations of pine marten and to guide the selection of potential reintroduction sites. Recently founded populations in southern England are small, fragmented and highly vulnerable to environmental, genetic and demographic factors (Birks et al., 2025, MacPherson et al., 2014; Gilpin and Soulé, 1986). Interventions, including translocation and reintroduction projects, are therefore necessary for the long-term viability of populations in Britain (MacPherson and Wright, 2021). Previous feasibility assessments for the reintroduction of pine marten to Britain have used occurrence records from populations in Scotland, Ireland and the Netherlands to predict habitat suitability in Britain at 1 km resolution (MacPherson et al., 2014; MacPherson et al., 2024; Scopes et al., 2025). Our methodology uses occurrence data from across the global range of pine marten, to predict suitability at 100 m resolution. Pine marten populations in the UK and Ireland have likely been driven to suboptimum, or ‘refuge’, areas due to anthropogenic disturbance, with loss of forest cover as a main contributor (European

Environment Agency, 2018a). Across mainland Europe, we assume that pine marten can more easily disperse and occupy their preferred habitat due to greater forest cover and connectedness. Consequently, we expect our model predictions, using global occurrences and a high resolution reflecting the habitat scale used by pine martens, to accurately identify areas of optimal habitat suitability. resolution capturing the habitat scale pine martens operate at, will reflect the optimal habitat suitability requirements of pine marten.

## Methods

We assessed pine marten habitat suitability at two distinct spatial scales and extents. First, we fit eSDMs across Europe at 1 km resolution to represent the species’ global distribution, before creating fine scale maps for Great Britian at 100 m resolution to better inform potential release sites for future reintroductions.

### Occurrence data

The occurrence data was collated from the Global Biodiversity Information Facility (GBIF, 2023), which holds a vast array of georeferenced species occurrence records, and the NBN Atlas (NBN, 2023), the UK’s main biodiversity data centre. GBIF provided 26,277 presence records across Europe (https://doi.org/10.15468/dd.32w6qw), and NBN Atlas provided 11,048 records across Britain and Ireland (Supplementary material 1). Although these databases hold large datasets, many records may be subject to bias from uneven sampling or inaccurate georeferencing (Glon et al., 2017; Beck et al., 2014). For example, certain areas may be the focus of targeted pine marten surveys (Austin, 2007).

Data cleaning is essential so that models are built on robust and spatially-uniform data. To clean the GBIF data we followed the methodology of Zizka et al (2019) and used the ‘CleanCoordinates’ package v2.0-11 (Zizka et al., 2019). This included removing records made prior to 1945 (which have low accuracy), removing duplicate records, removing coordinates assigned to country and province centroids, removing records with 0,0 coordinates, coordinates at sea, and coordinates assigned to biodiversity institutions. We incorporated additional cleaning steps to the remaining data including removing records with coordinate uncertainty above 1 km, removing records of unknown source, records derived from fossil specimens, and records with noted geospatial issues. After data cleaning, 32,000 out of 37,325 records were retained (Fig. 1).

**Fig 1.**
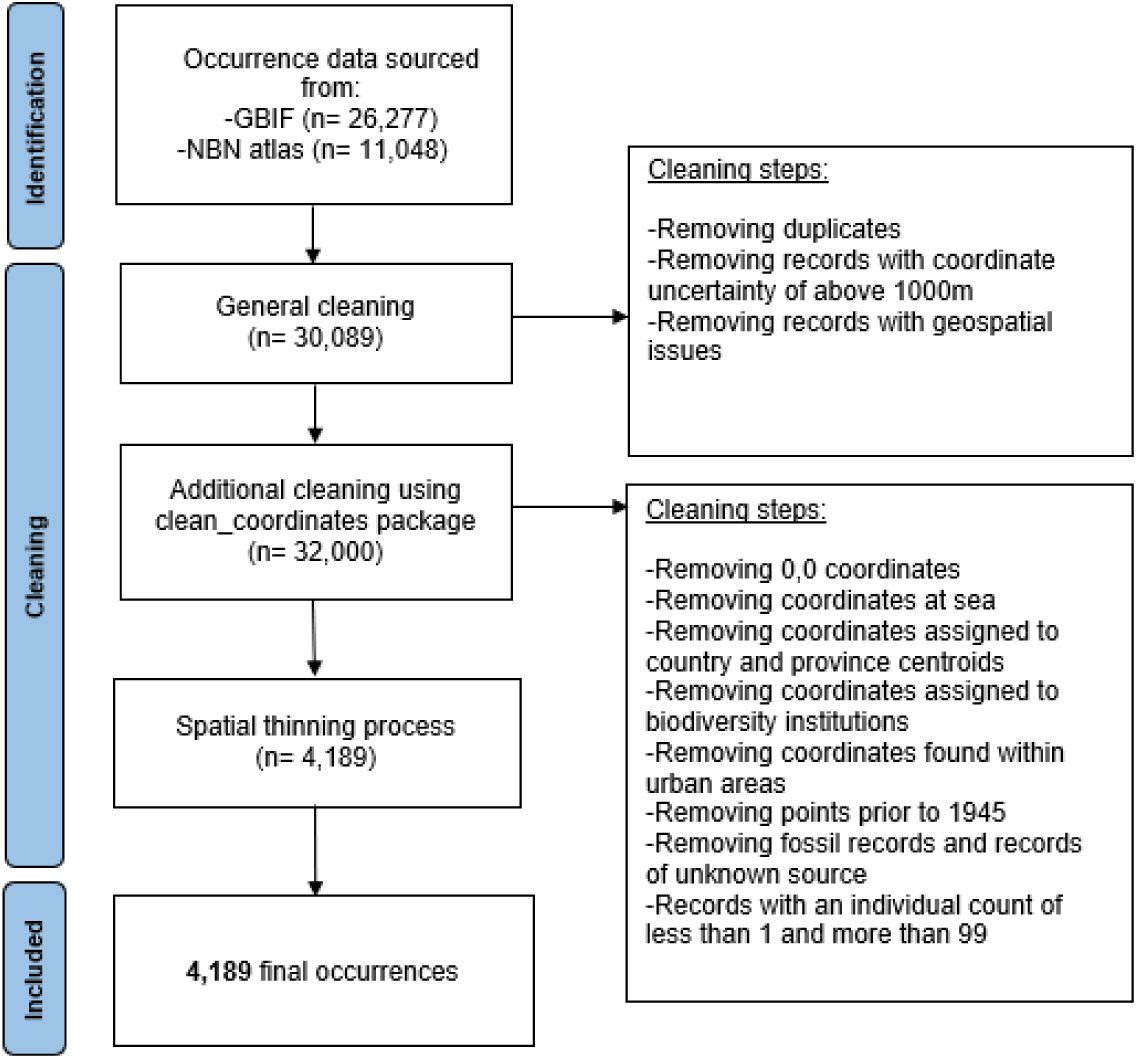
PRISMA diagram illustrating the necessary cleaning steps with the number (n) of occurrences remaining after each cleaning process.

Sampling bias can skew model accuracy, as it may not reflect the complete species’ range of environmental conditions and may over-represent environmental conditions associated with regions of higher sampling (Kadmon et al., 2004; Anderson and Gonzalez, 2011; Duque-Lazo et al., 2016). To reduce the influence of spatial bias, occurrence data should undergo spatial thinning. To do so, the ‘spThin’ function in R was applied to the cleaned occurrences, which used a randomised approach to remove occurrence points within 10 km of one another, making sure the maximum number of occurrence points were returned (Aiello-Lammens et al., 2015).

We identified 10 km as an appropriate thinning distance by visually inspecting a variogram of the environmental variables (Fig. 2). The variogram shows the variation in environmental conditions with increasing Euclidian distance between points. Therefore, the distance at which the variance ceases to increase and begins to plateau can inform the distance at which occurrences are geospatially independent from one another, and therefore the distance to use for thinning (Anderson and Gonzalez, 2011; Bio et al., 2002). After data cleaning and spatial thinning, occurrences were clipped to the model area of Europe, retaining 4,189 occurrences for use in the modelling (Fig. 3).

**Fig 2.**
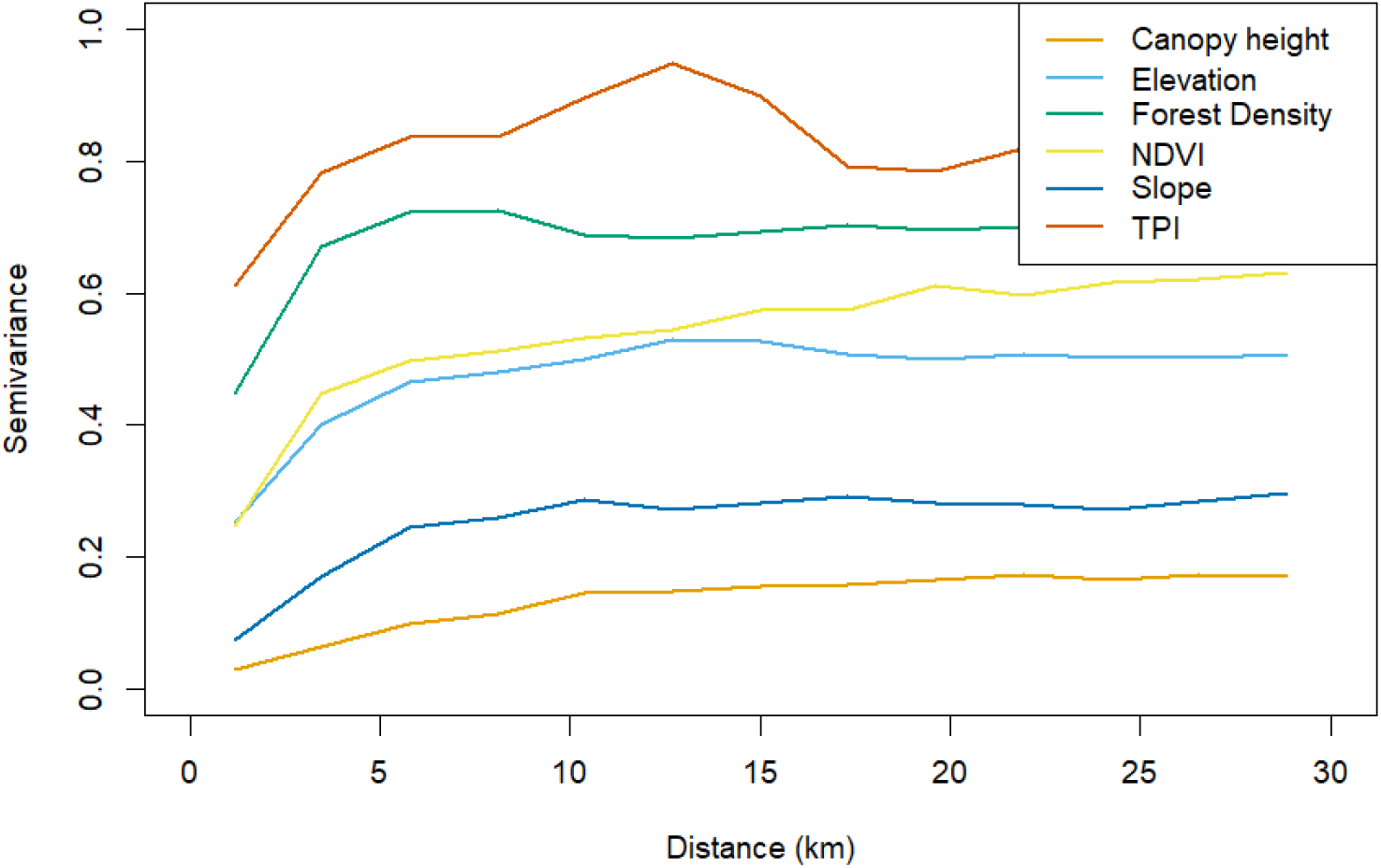
The variogram shows that the semivariance reached a plateau after 10 km from points. This ‘practical range’ was used to thin the occurrence data in order to remove spatial autocorrelation, but retain the most occurrence data.

**Fig 3.**
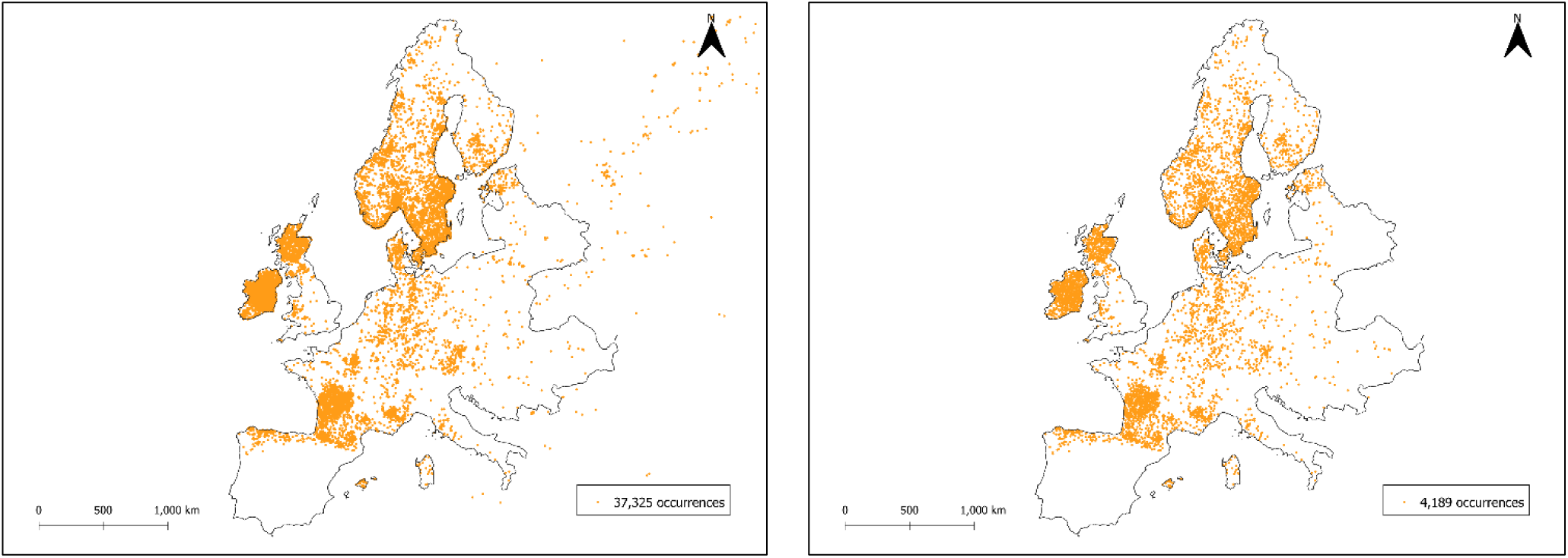
The locations of occurrence records before (left) and after (right) cleaning and spatial thinning within the modelled area.

### Environmental predictors

SDMs are based on the ecological niche concept, which posits that species occupy spatial ranges with suitable environmental conditions for their existence. Thus, the environmental variables used to predict suitability herein were chosen based on the ecological requirements of pine marten. It is hypothesised that the suitability of habitat for pine marten is determined by risks of mortality, food availability, and availability of denning sites (Stringer et al., 2018; MacPherson et al., 2014). A total of fourteen environmental variables were chosen based on these factors. Processing of environmental predictors was conducted in QGIS v3.22 (QGIS.org, 2021). All environmental variables were reprojected to ETRS89-extended /LAEA Europe, resampled to a resolution of 1 km (nearest neighbour for categorical variables and bilinear for continuous variables using WARP tool in QGIS), cropped to the same extent of Europe and aligned. Four of these environmental variables were used for the British projection, whilst the remainder were sourced from different datasets (Table 1).

**Table 1:**
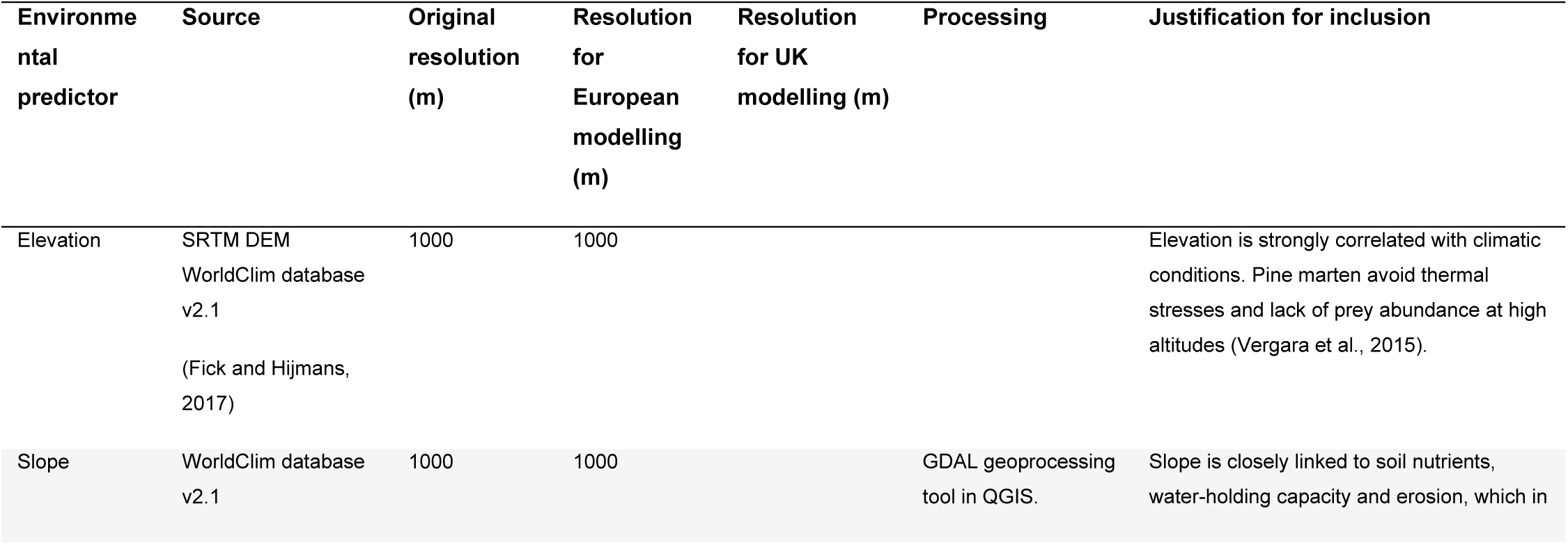
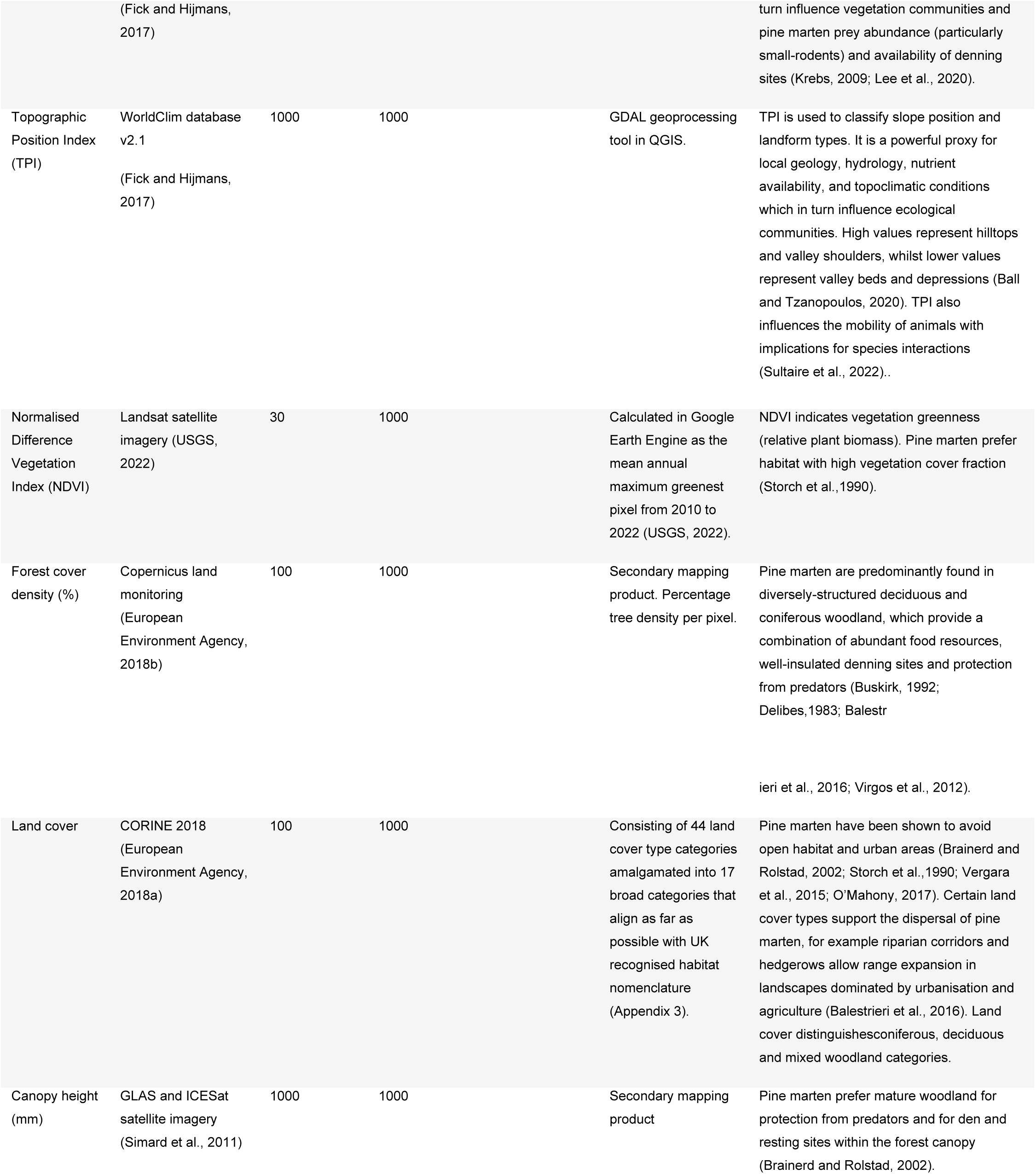
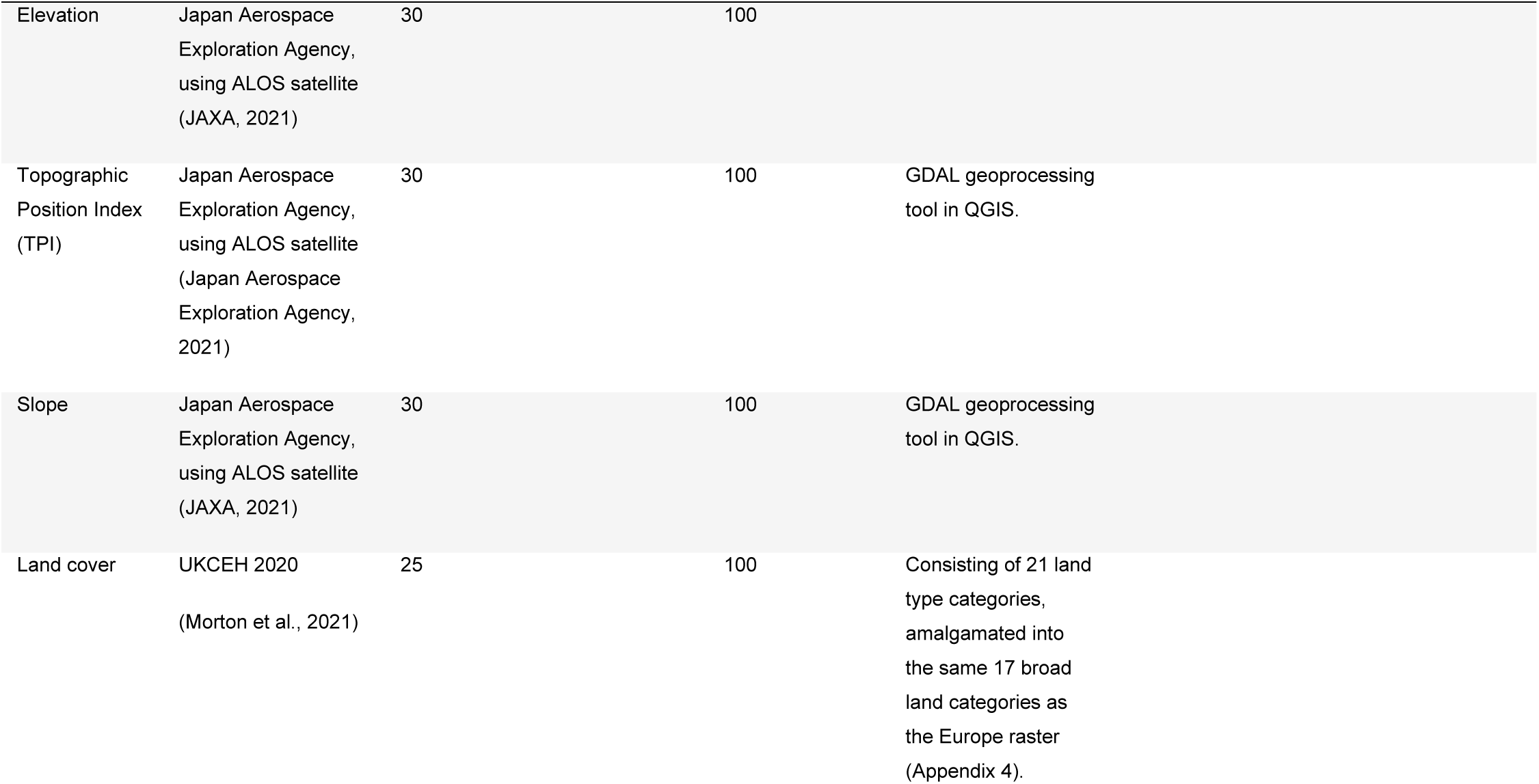
Environmental predictors used in the modelling or removed after multicollinearity assessment, with their justification for inclusion based on ecological relevance.

Multi-collinearity was tested between environmental variables using Pearson’s r-correlation test with r >0.7 considered as unacceptably collinear. Collinear variables cannot be included in statistical models because it would be impossible to determine the unique effect of each variable (Dormann et al., 2013; Brun et al., 2019; Glon et al., 2017). This process led to the removal of five environmental variables: average monthly maximum and minimum temperature (°C) derived from WorldClim database v2.1 (Fick and Hijmans, 2017), average monthly precipitation (mm) derived from WorldClim database v2.1 (Fick and Hijmans, 2017), and northness and eastness derived from SRTM WorldClim database v2.1 (Fick and Hijmans, 2017). Removal of these variables was not a concern as average elevation, collinear to climatic variables, was retained, whilst northness and eastness are unlikely to influence pine marten habitat suitability.

Two environmental variables were removed from the eSDM due to spatial autocorrelation of sampling efforts (sampling bias); distance to roads and urban areas. Distance to roads was derived from a shapefile subset to major roads (A and B roads) using UN Spatial Data Infrastructure Transport (UNSDI-T) v2 (CIESIN, 2013). Distance to urban areas was derived from a shapefile from North American Cartographic Information Society (Patterson et al., 2012). Both urban areas and major roads vector layers were rasterized, and distance was calculated from the proximity (raster distance) tool, both derived from GDAL in QGIS.

This process resulted in a total of seven environmental variables used to inform suitability in the model (Table 1): elevation, slope, topographic position index, normalised difference vegetation index (NDVI), forest density, land cover, canopy height.

### Modelling

#### Model algorithms

To build the ensemble model, we used 9 algorithms within the BIOMOD2 v 4.2 package in R (R Core Development Team, 2014; Thuiller et al., 2016).These were generalised linear model (GLM), generalised boosted regression model (GBM), generalised additive model (GAM), classification tree analysis (CTA), artificial neural networks (ANN), flexible discriminant analysis (FDA), multivariate adaptive regression splines (MARS), random forest (RF), and maximum entropy model (MaxEnt), using the default parameters specific to each algorithm.

#### Pseudo absences

Most SDM algorithms require both presence and absence data. For many species, true absence data is rarely available or unreliable and so presence-only models are used, wherein artificial absence data, known as pseudo-absences, are generated (Barbet-Massin et al., 2012). Pseudo-absences aid the model’s ability to differentiate the environmental conditions under which the species occurs (Brown and Yoder, 2015). We used a large number of pseudo absences, equally weighted to presences (4,189), as has been shown to produce the most accurate predictions (Barbet-Massin et al., 2012). Pseudo-absences were randomly generated within the entire study area. We used BIOMOD’s disk method to prohibit pseudo-absence generation within 5 km of occurrence points, respective of a typical pine marten home range for females (Stringer et al., 2018). This random assignment of geographically stratified pseudo-absences is recommended for machine-learning and classification methods (Barbet-Massin et al., 2012). To reduce bias towards a single distribution of pseudo-absences, five sets were generated (Guisan et al., 2017). Prevalence was assigned as 0.5, where presences and pseudo-absences have the same weighting in the model calibration process (Liu et al., 2005; McPherson et al., 2004).

#### Model evaluation

To test model accuracy when no independent data is available, cross-validation is performed, where the data is randomly partitioned into a set of training data and testing data (Raes and Aguirre-Gutiérrez, 2018). For our analysis, data was randomly partitioned to 70% for training and 30% for testing using repeated split samples. This cross-validation process was repeated 10 times for each modelling run (Yu et al., 2024). We chose the widely used repeated random cross-validation method. Model quality was evaluated based on kappa statistic (Cohen, 1960), the area under the receiver operating characteristic curve (AUC), and the true skill statistic (TSS) (Allouche, et al., 2006). Only single models with a TSS score greater than the upper quartile (0.324) of TSS scores (0%: 0.218, 25%: 0.286, 50%: 0.307, 75%: 0.324, 100%: 0.368) were retained for the final ensemble and ensemble projections (Tanaka et al., 2020; Hill et al., 2017). TSS corresponds to the sum of sensitivity and specificity minus one (Barbet-Massin et al., 2012). TSS scores can be classified according to Ben Rais Lasram et al., (2010) as excellent (TSS > 0.8), good (TSS = 0.6–0.8), fair (TSS = 0.4–0.6), poor (TSS = 0.2–0.4), and no predictive ability (TSS < 0.2). We chose TSS to evaluate model accuracy for inclusion in the ensemble models as it is a threshold-dependent measure of model accuracy and independent of prevalence (the ratio of presence to pseudo-absence data in the presence-absence predictions) (Allouche et al., 2006).

We combined the single models into ensemble models based on mean, median and weighted average. Mean and median produce the ensemble results by averaging or taking the median of the single model probability of suitability values, respectively, whereas weighted mean weights the individual models based on their cross-validation performance, in our case the TSS values (Marmion et al., 2009; Thuiller et al., 2009). For the ensemble model evaluation, the same evaluation statistics from the single models were used to assess model performance. The method used to combine the models that performed best, should be used to present the projection results. The ensemble model included 111 single models out of 450 individual models.

To evaluate the relative importance of each environmental variable, a simple Pearson’s correlation *r* was calculated between predictions of the full model and a model with each variable removed (Thuiller et al., 2009). We also generated species response curves, which show how suitability for pine marten changes as a given environmental variable changes.

#### Habitat suitability projections

We generated single and weighted mean ensemble model projections in Britain using the 100 m resolution environmental rasters. Suitability index has been re-scaled to a 0 to 100 range, where larger values indicate higher suitability.

## Results

Our study used eSDM with 4,189 pine marten occurrences from across the species’ range and 7 environmental variables to predict pine marten habitat suitability in Europe and Great Britain. The results indicate that elevation, land cover and vegetation greenness (NDVI) are the strongest predictors of habitat suitability. Across the extent of Europe at 1km and Britain at 100m, the eSDM results illustrate high suitability for pine marten in areas with woody vegetation cover in low topographic positions, and notably low suitability in urban areas and extensive areas of arable land.

The mapped relative probability of habitat suitability in Europe on a 1km scale shows markedly low habitat suitability for pine marten at high altitudes in the Alps and the Scandinavian Mountains, in heavily modified and farmed landscapes, in dry and arid zones including much of Spain, and over open water bodies. Higher habitat suitability occurs in areas with woody vegetation cover, in lowlands, low topographic positions, and notably along Europe’s West coast (Fig. 4).

**Fig 4.**
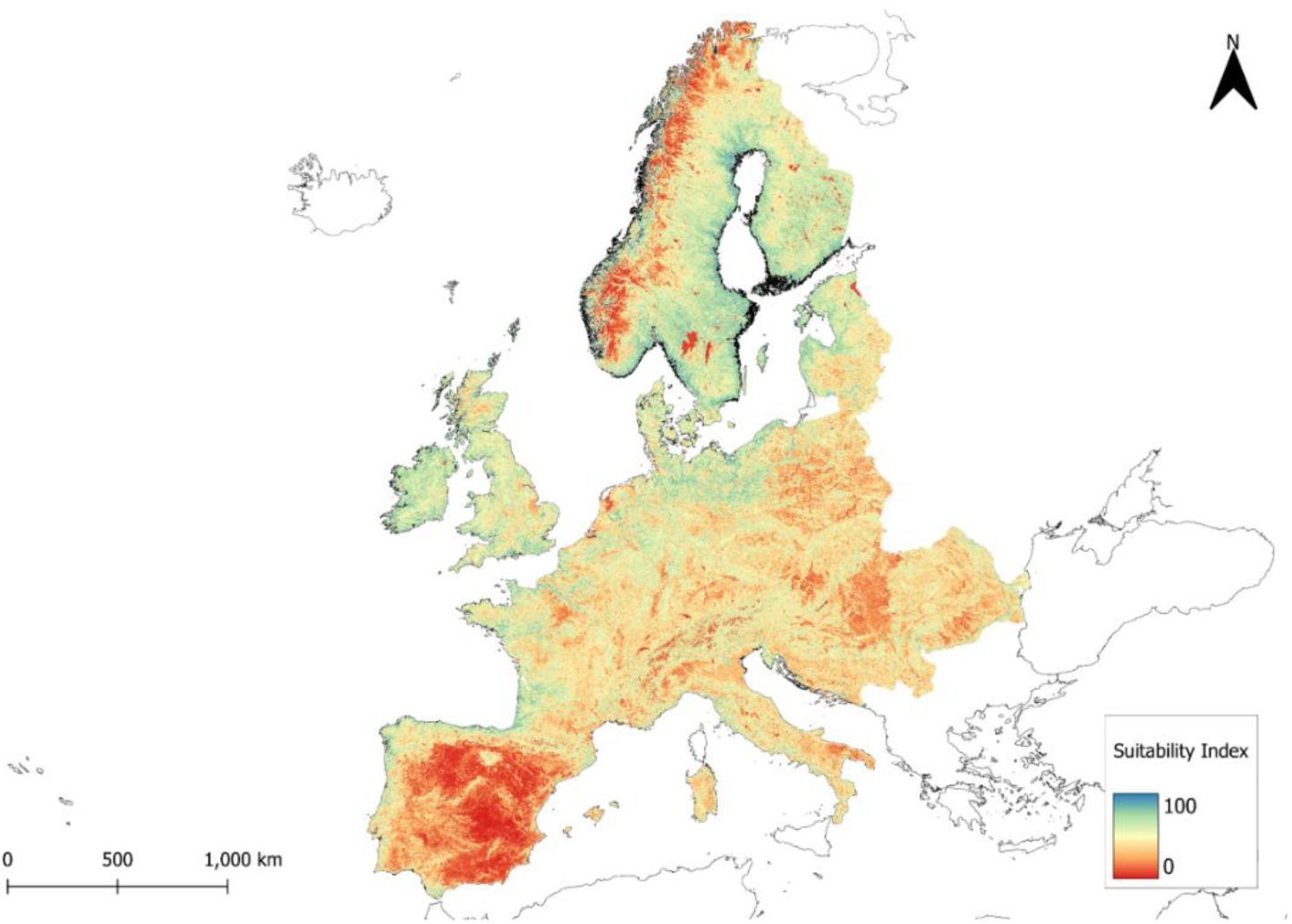
Habitat suitability map indicating pine marten suitability across Europe at a resolution of 1km x 1km. Suitability index ranged from 0 to 100, where larger values indicate higher suitability.

The mapped relative probability of habitat suitability in Great Britain on a 100m scale shows high suitability across substantial areas in the South East of England, South West of England, East Yorkshire, and Gloucestershire, with pockets of suitable habitat along the West Coast of Britain. Suitability is greatest in wooded landscapes and lower in urban and intensively farmed areas. Substantial areas of the Scottish Highlands, North Pennines, Central Midlands and South Lincolnshire show relatively low suitability (Fig. 5).

**Fig 5.**
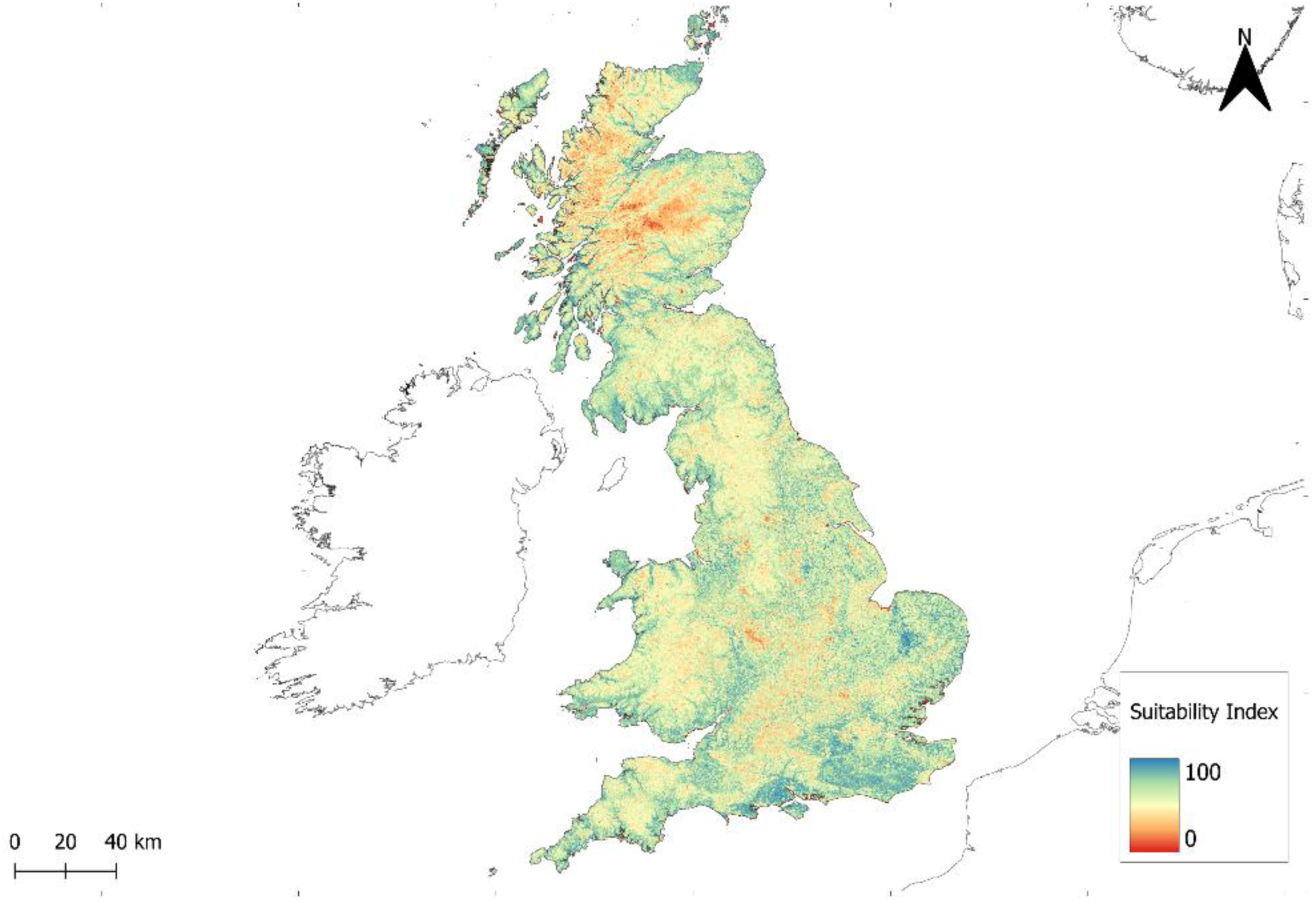
Habitat suitability map indicating pine marten suitability across Great Britain at a resolution of 100 × 100 m. Suitability index ranged from 0 to 100, where larger values indicate higher suitability.

TSS and ROC validation scores were used to assess the performance of the models. The mean TSS value of the single models was 0.3046 (Min: 0.218, Q1: 0.286, Median: 0.307, Q3: 0.324, Max: 0.368). The mean ROC value of the single models was 0.7087 (Min: 0.628, Q1: 0.698, Median: 0.713, Q3: 0.723, Max: 0.756). The calibration scores for the mean ensemble model were TSS = 0.425 and ROC = 0.786. The calibration scores for the median ensemble model were TSS = 0.361 and ROC = 0.751. The calibration scores for the weighted-mean ensemble model were TSS = 0.426 and ROC = 0.786.

The ensemble model variable importance results, averaged across the ensemble types, showed elevation was the strongest predictor of suitability, as were land cover, vegetation greenness (NDVI) and slope but to a lesser degree. TPI and canopy height were of relatively low importance, with forest density being of least importance (Fig. 6).

**Fig 6.**
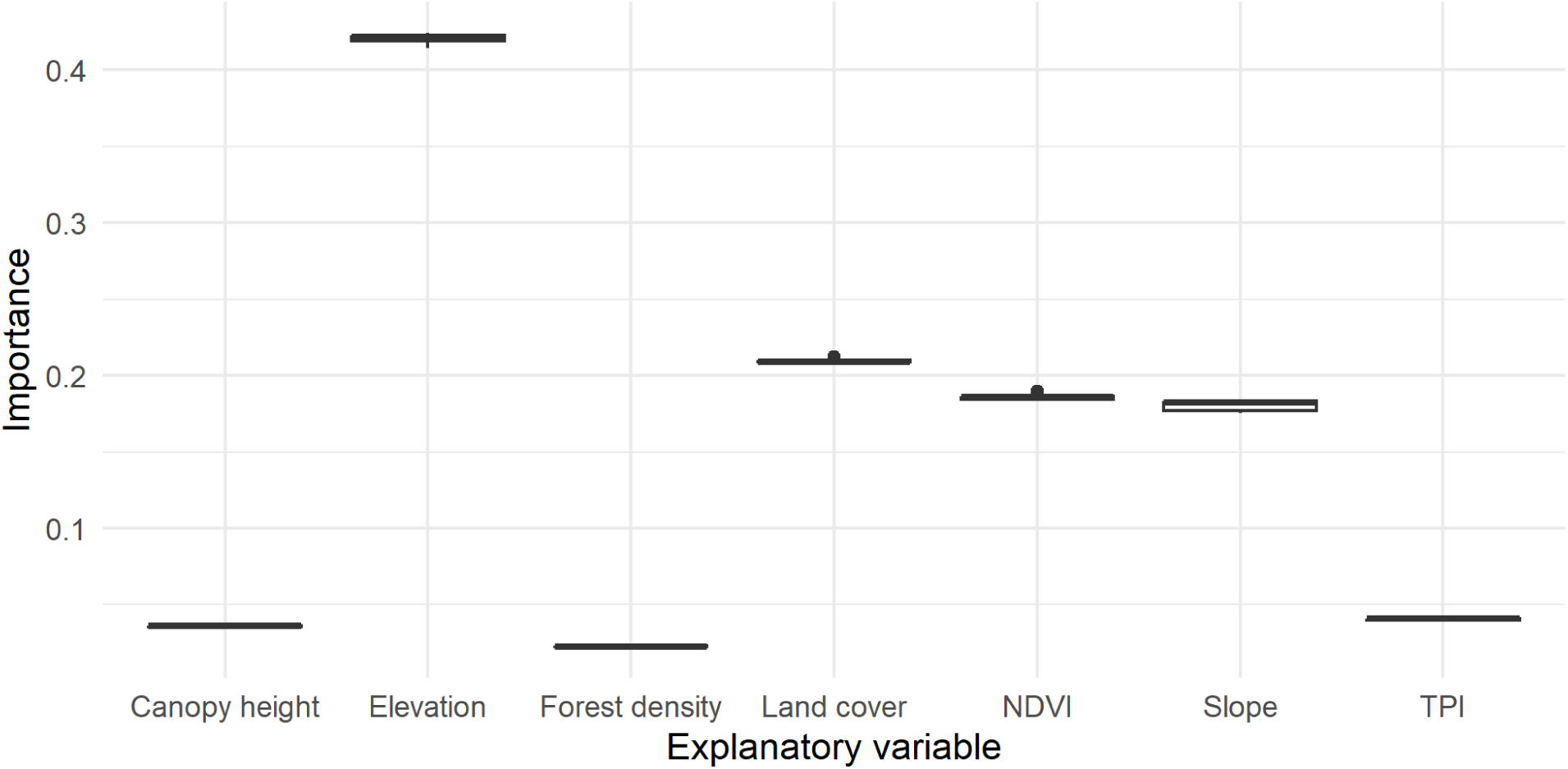
Box plot showing the ensemble variable importance, across the mean, median and weighted mean.

The response curves, averaged across the ensemble results (Fig. 7), indicate that pine marten favour lower elevations, with a sharp decline in suitability at elevations above 500 m above sea level (a.s.l.).

**Fig 7.**
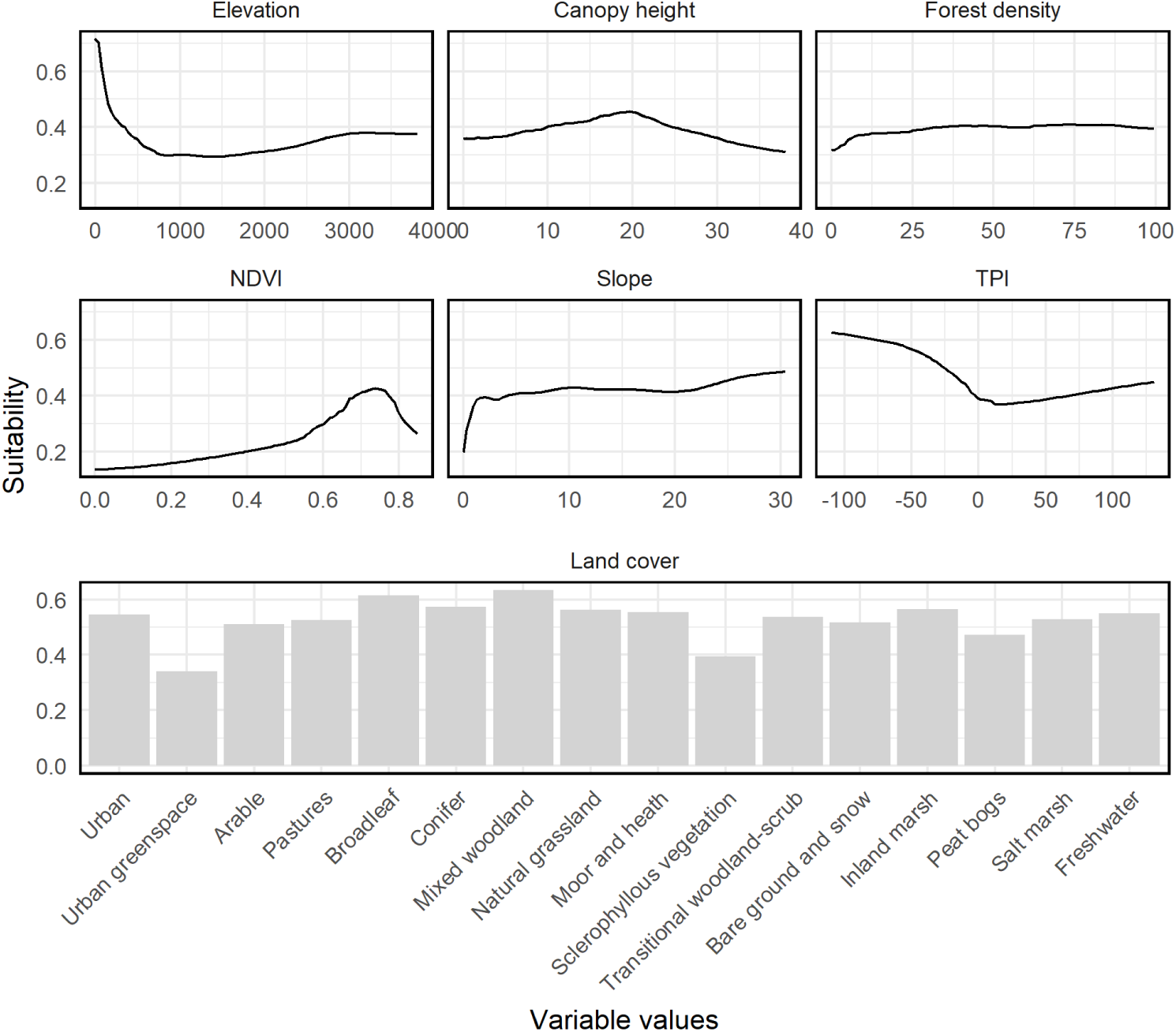
The response curves, averaged from the final ensemble models. Probability of suitability is on a 0-1 scale.

Woodland was favoured over other land cover types, with mixed woodland as the most suitable, followed by broadleaf woodland and coniferous woodland (Fig. 7). Preference for arable and pasture land cover was lower, with suitability values below 0.5. Urban greenspace, sclerophyllous vegetation and peat bogs were the least favoured land cover type.

There is a fairly consistent medium probability of occurrence of pine marten across all values of tree density, although at low densities (<25% tree cover) probability starts to decrease, and at very low densities (<6% tree cover), the probability of occurrence drops sharply from about 0.42 to 0.36 (Fig. 7). NDVI showed a peak in suitability at 0.75 NDVI, where the probability of occurrence exceeds 0.4 (Fig. 7) and drops suddenly to 0.25 suitability probability. The probability of occurrence of pine marten increases with canopy height, until canopy height reaches above 20 m and probability of occurrence drops from 0.4 at 25 m, to 0.3 at 35 m (Fig.7).

The species response curve for TPI, indicates a preference of pine marten for lower topographic positions within valleys and depressions (Fig. 7). The probability of occurrence of pine marten is fairly consistent across all values of slope, apart from a slight decrease in very flat areas (Fig.7).

## Discussion

In this study, eSDMs were used to map environmental suitability for pine marten in Great Britain, to inform conservation and restoration of the species, with a focus on reintroduction site suitability. Across the extent of Europe, the 1km eSDM results indicate high suitability in the British isles, northern parts of continental Europe including France, Denmark and Germany and lower altitudes of Scandinavia. Low suitability regions were often found at lower latitudes including the Mediterranean regions such as large parts of the Iberian Peninsula and South East Italy, as well as areas of high altitudes including the Alps, the North Western Highlands across Scandinavia and extents of the Scottish Highlands (Fig. 4). Across Britain, the 100m eSDM results indicated high suitability in most regions of the South East of England, parts of the South West of England, East Yorkshire, Gloucestershire and eastern borders of East Anglia, with pockets of suitable habitat on the West coast of England (Fig. 5). High suitability identified in the South East of England could help to inform potential release areas of the South East Pine Marten Restoration project, who are currently scoping a reintroduction to the High Weald National Landscape. Here, we found areas of high suitability to be extensively wooded environments in protected areas (National Landscapes and NPs), with lowest suitability in urban and especially low-lying coastal urban areas (Fig. 6). Notable hotspots of suitability were in the South Downs National Park, the High Weald National Landscape, and the western end of the Kent Downs National Landscape which may serve as suitable reintroduction locations.

Model results show low suitability at high altitudes, particularly in the Scottish Highland regions. Significant distributions of pine marten across Scotland does not necessarily correspond to high-quality habitat, rather, it may represent remnant or refuge areas where populations persist in sub-optimal habitats in Britain. By incorporating European occurrence data to build our eSDMs, the model provides a more ecologically representative assessment of habitat suitability for the species, and one which is relevant to expected poleward range shifts under future climatic conditions.

The variable importance results showed that elevation is the strongest predictor of habitat suitability for pine marten across its global range. Our results show substantially higher probability of pine marten habitat suitability at lower elevations (< 500 m a.s.l). This preference for lowland wooded habitats in Europe is likely to be linked to the availability of denning sites which is higher in mature broadleaved woodland which typically occurs at lower elevations (Harmer et al., 2010; Portoghesi, 2006). Coniferous woodlands and alpine tundra and meadows generally occur at higher altitudes but lack the abundance of prey and denning sites required by pine marten (Fonda et al., 2021). Based on our results we see that pine marten are unlikely to occur in substantial populations at altitudes above 2000 m a.s.l.

Although its relative importance was low, the response curve indicates that pine marten prefer lower topographic positions, representing areas within valleys and depressions. We expect this preference is linked to the greater presence of woody vegetation in these locations compared to sparser tree cover that tends to occur in higher topographic positions, on ridges and valley shoulders. This can be seen in the South East of England where areas with highest suitability for pine marten are in topographically complex, often forested, areas (Appendix 1).

Land cover is the second strongest predictor of suitability, with mixed, broadleaf and coniferous woodland being highly favoured (Fig. 7). As forest-dwellers, pine marten are found primarily in well-structured deciduous and coniferous woodland, with well-insulated denning sites and abundant food resources, and with a complex understorey providing a low risk of predation (Buskirk, 1992, Virgo et al., 2012; Delibes,1983; Balestrieri et al., 2015). The results show a lower preference for coniferous woodland compared to mixed and broadleaf woodland, which could reflect their preference towards more structurally diverse woodland systems and lower canopy heights associated with avoidance to coniferous-dominant plantation habitat. Planted coniferous woodland is often thinned and structurally uniform, which serves as a poor habitat for pine marten due to the lack of diverse ground flora, foraging opportunities and denning sites.

Interestingly, arable, pasture and grassland land cover types were found to be moderately favoured, contradicting previous models which have shown that they avoid open habitats (Brainerd and Rolstad, 2002; Storch et al.,1990; Vergara et al., 2015; Wereszczuk and Zalewski, 2015; O’Mahony, 2017). This may be explained by the species’ ecological flexibility and ability to occupy heterogenous landscapes. Evidence suggests that diverse habitats can promote higher diversity of food resources than a continuous forest habitat (Rosalino and Santos-Reis, 2009; Vergara et al., 2015). For instance, we know that pine marten travel up to 150 m from woodland edge to forage for prey species such as field voles in their favoured habitats, provided there is some low-level vegetation cover (Caryl, 2008).

Linear or small habitat features will be underrepresented in our model due to the analytical resolution of the modelling across both projections (1km and 100m). For example, shrub or hedge features within a grassland or arable pixel will not be represented. These features can serve as important facilitators of connectivity for individuals to move through the landscape to access other patches of woodland, therefore our model likely underpredicts habitat suitability. Notably, urban greenspace, sclerophyllous vegetation and peat bog land cover types were least suitable. These land types are also rare. If repeating this study, one may consider further reducing the total number of land cover types by conducting pairwise correlations between the broad land cover types to combine correlated categories to simplify the model (Buckley and Lundy, 2012).

Vegetation greenness (NDVI) was the third strongest predictor of suitability because pine marten inhabit woodland and adjacent edge habitat. As well as providing denning sites and shelter, vegetation greenness is also strongly correlated with prey abundance. For example, rodent density is strongly correlated with net productivity of ground vegetation (Zalewski et al., 2004). The response curve showed a peak in suitability around 0.55-0.8 NDVI which indicates high vegetation cover fraction like that of tall grassland and woodland canopy.

Our results also suggested that pine marten will occupy a broad range of forest densities, with a consistent medium probability of occurrence of pine marten across all values of tree density (Fig. 7). We expect this links strongly with their ability to adapt to possibly sub-optimum environments (MacPherson et al., 2014). For example, Southern European populations can be found across ancient woodland, coppiced woodland and cultivated land with forest fragments, where they are not restricted to dense woodland cover (Pittiglio, 1996). Even within countries, the adaptation to varying vegetation structures is evident. For example, French populations have genetically diverged to occupy varying woodland cover. In the Bressie region, individuals occupy human-dominated landscapes with fragmented forest whilst in the Ariège region individuals occupy densely forested habitat (Larroque et al., 2015).

The results show canopy height was a relatively unimportant variable, however suitability increased with increasing canopy height, up to 20 m, after which it decreased with increasing canopy height. Pine marten favour mature forest for protection from predators and access to denning and resting sites within the forest canopy (Brainerd and Rolstad, 2002).. However, the decrease in suitability in very tall forests (>20 m) may be due to fewer denning sites in less structurally diverse, coniferous-dominant plantation habitats (Croose et al., 2016).

Mean monthly temperature (°C) and mean monthly precipitation (mm) were excluded from the eSDM. Although these variables have strong predictive power at the European scale, the climate of Great Britain is already highly suitable for pine marten. Consequently, including broad-scale climatic predictors led to their undue dominance in the model and reduced its ability to map fine-scale habitat suitability within Great Britain. In a previous model iteration with these variables included, mean monthly temperature accounted for 37% of model influence and mean monthly precipitation for 17%, making them the first and third most important predictors, respectively. Model evaluation metrics were also slightly higher when these variables were included (mean single-model validation TSS ≈ 0.43). However, mean monthly precipitation showed collinearity with vegetation greenness (NDVI) and elevation, further justifying its removal.Sampling bias is a major challenge for species distribution modelling, and we identified disproportionate sampling effort in our dataset. Specifically, pine marten occurrence records were more common near urbanised areas and roads, possibly linked to accessibility for recorders (Araújo and Guisan, 2006). This could lead to spatial bias in the model, misrepresenting the environmental conditions which reflect suitability. We therefore decided to remove the distance to roads and urban areas variables to prevent the influence of spatial autocorrelation. One may also argue that roads and urban areas should not be included as they serve as barriers rather than predictors of suitable habitat. In other words, pine marten do not deliberately choose to be close to or far from roads, rather, territory is chosen based on predominantly the abundance of woodland for denning potential and availability of prey. The impacts of roads and urban areas on habitat suitability and connectivity are better examined using connectivity modelling to quantify the effects of movement barriers, which includes analysing the risks of mortality and how it effects overall population viability. Whilst comprehensive efforts have been made to reduce bias in the occurrence data, other causes of bias may still be present. For instance, we identified low environmental suitability at high altitudes above 2000 m, which may be due to a lower number of observations in these difficult to reach locations.

Our models did not perform to a good or excellent standard according to Kappa, TSS and AUC evaluation scores, which may be due to the large spatial extent of our analysis (Europe), which can lead to the selection of a higher proportion of less informative background points (Barbet-Massin et al., 2012). This can lead to difficulties detecting commission and omission errors (false presences and false absences), leading to low-moderate model performance. Moreover, climatic predictor variables were removed due to multi-collinearity, which had the strongest predictive power. Due to these limitations, the suitability maps should be used to guide further ecological feasibility research to assess suitability in the local context.

Raster data with higher resolutions than is typical for this type of study were used for a number of reasons. Firstly, higher resolution environmental variables provide more accurate information about the environment, whereas down-sampled environmental variables average across neighbouring cells, often leading to a poor measure of local conditions. This is particularly pertinent in highly fragmented landscapes, where environmental predictors that change over small spatial distances may become meaningless if a coarse resolution averages a broad range of values. Secondly, high-resolution projections provide suitability estimates for smaller land areas, which is particularly important in highly fragmented landscapes such as in England.

Producing eSDM maps at 100m resolution, improves our ability to inform potential reintroduction areas at a local scale.

To inform potential reintroduction locations, these results must be coupled with further research into barriers to movement, such as infrastructure, to inform the viability of a reintroduction.

MacPherson and Wright’s (2021) ‘Long-term strategic recovery plan for pine martens in Britain’ report recommends further research into the effects of roads on risk of mortality in more densely-populated parts of the Great Britain. Since male pine martens hold large territories, ranging from 2 km^2^ to up to 25 km^2^ (Zalewski and Jędrzejewski, 2006; Balestrieri et al., 2010), the requirement for connectivity between identified highly suitable habitat areas is essential for their long-term establishment (McNicol et al., 2020) and should therefore also be investigated. Connectivity modelling can also assess proximity to existing populations which is important for ensuring genetic diversity and the long-term viability of any reintroduced populations.

## Supporting information

R Script

## Appendices

## Appendix 1

Habitat suitability map indicating pine marten suitability across regions of Scotland, England and Wales at a resolution of 100 × 100 m. Suitability index ranged from 0 to 100, where larger values indicate higher suitability.

<u>North Scotland</u>

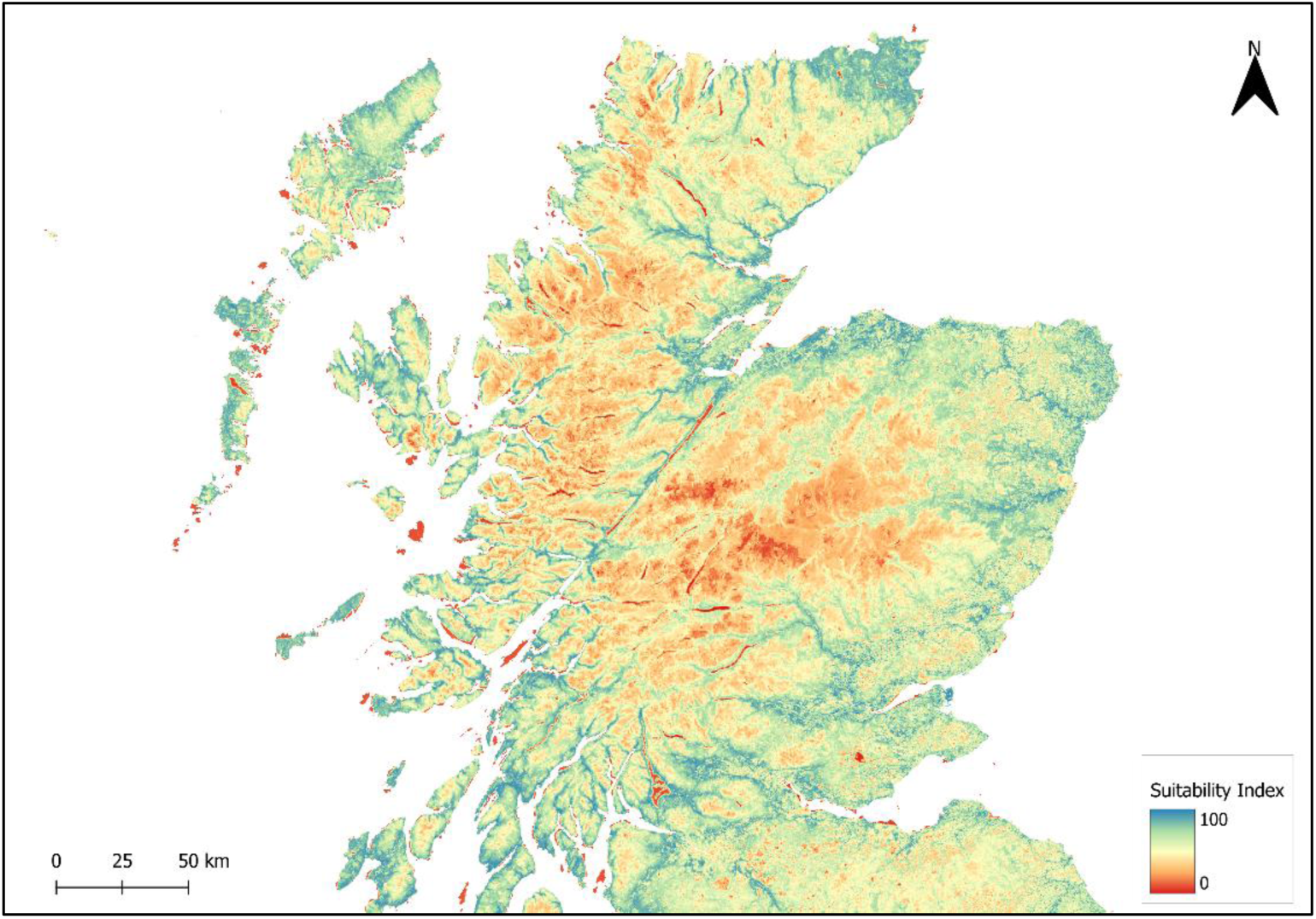

<u>South Scotland</u>

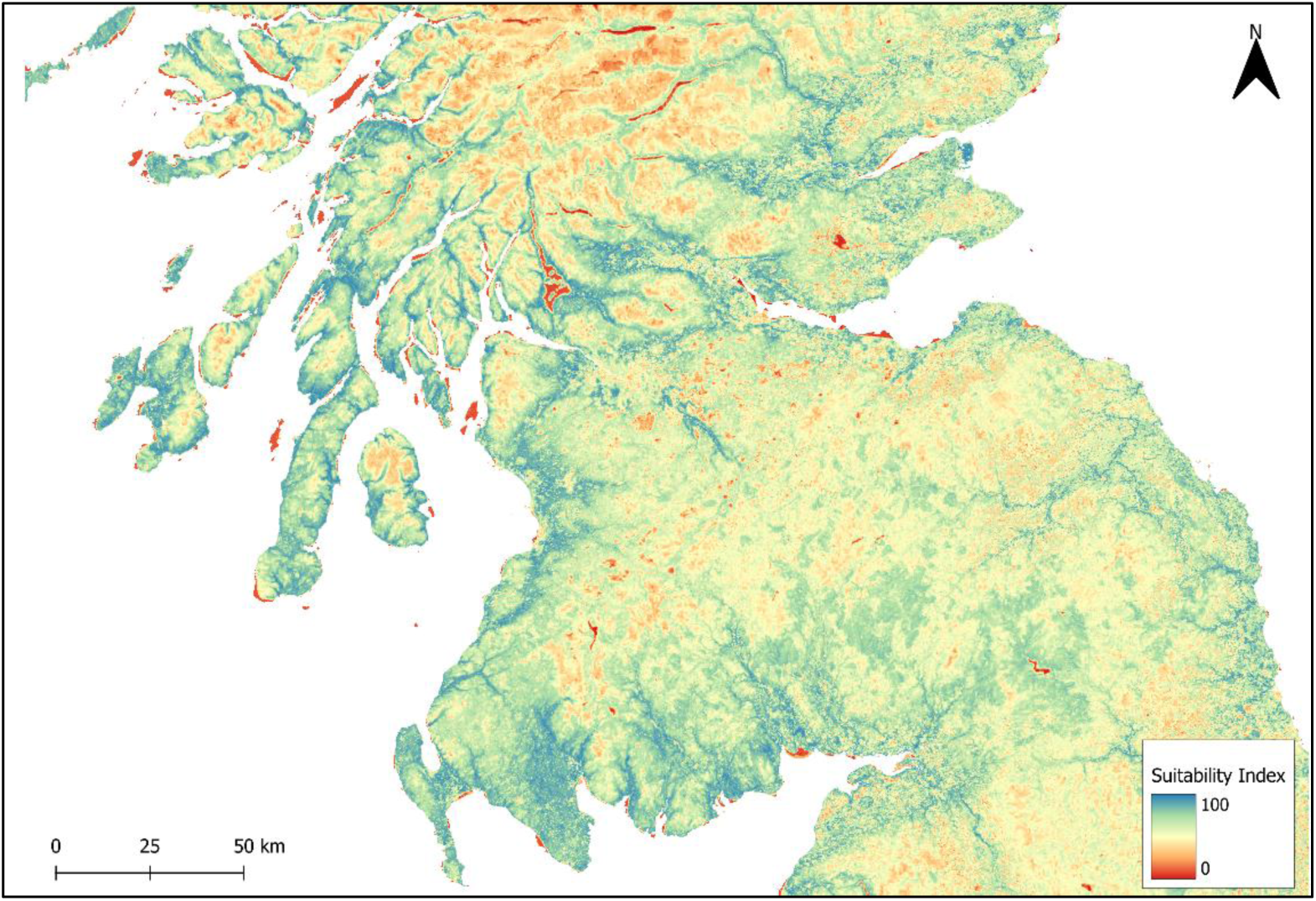

<u>Northern England</u>

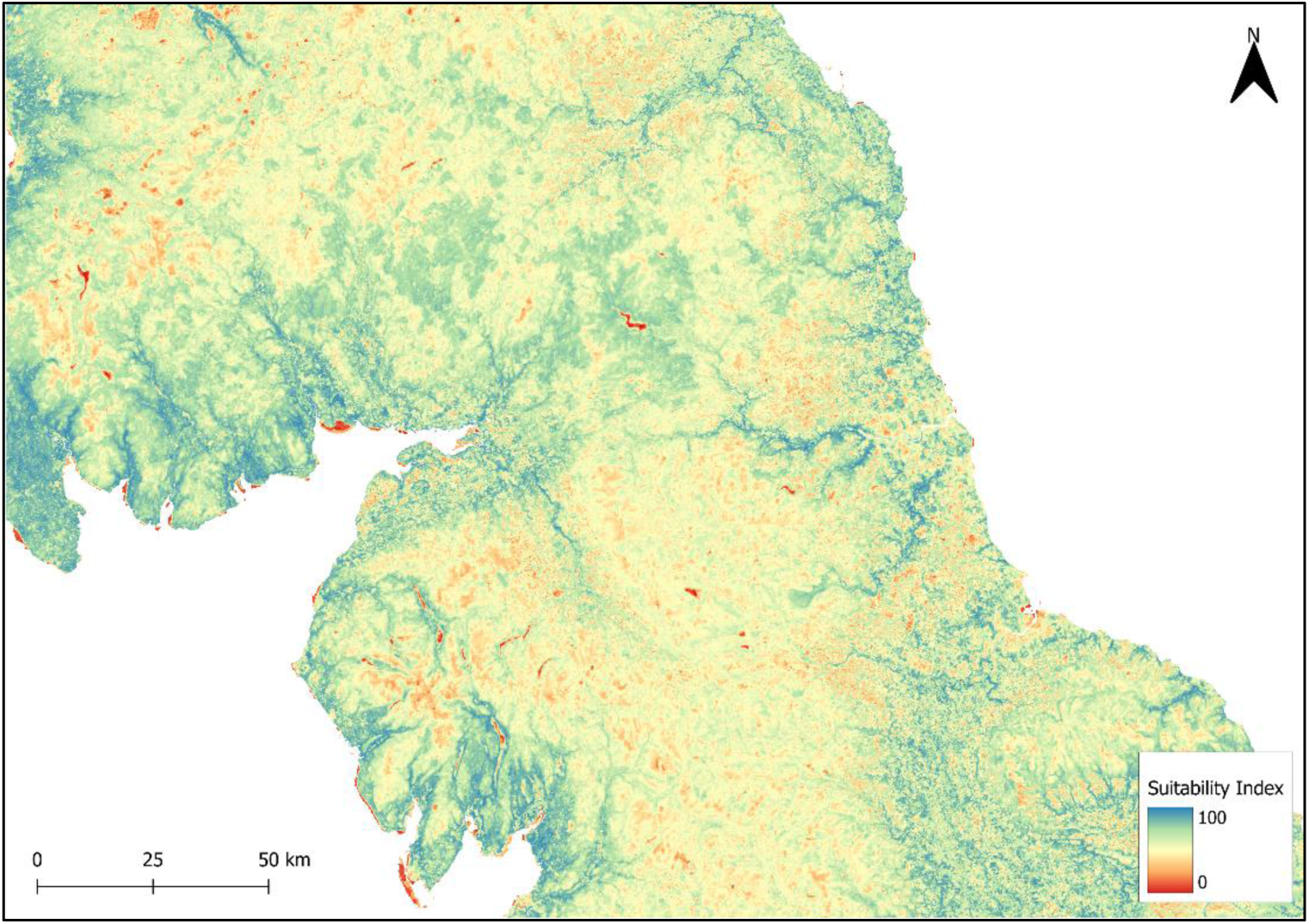

<u>Yorkshire</u>

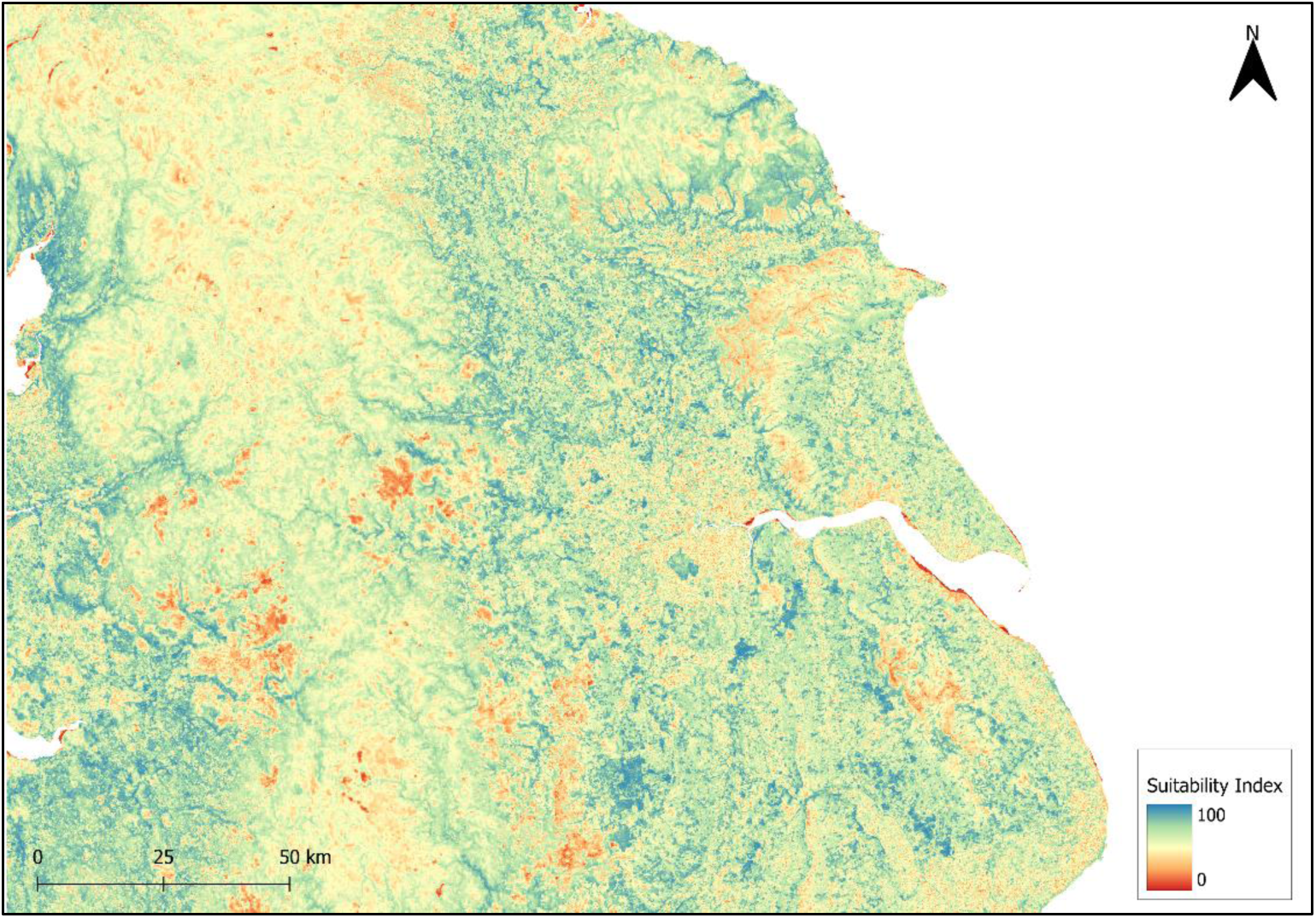

<u>Central England</u>

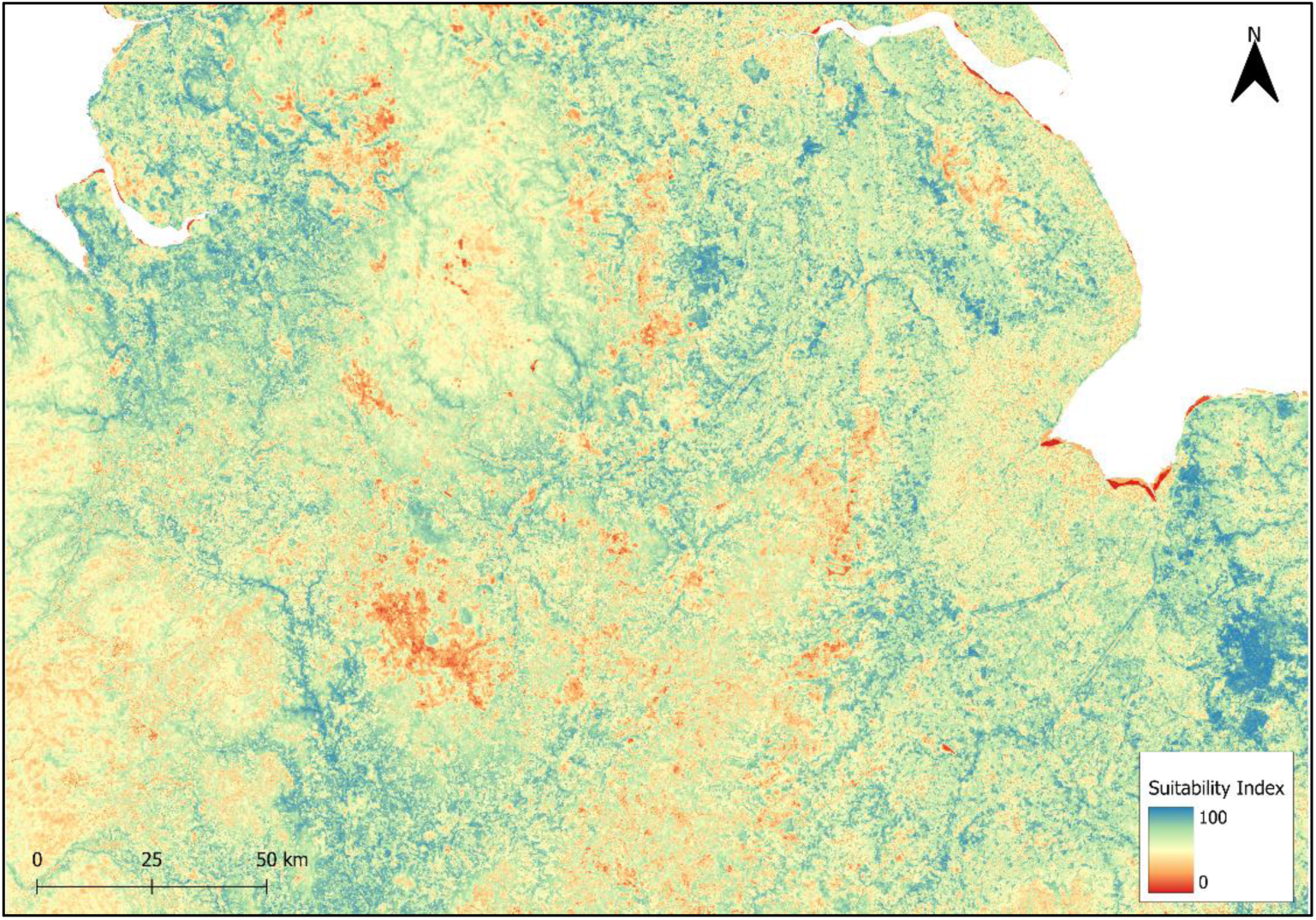

<u>Wales</u>

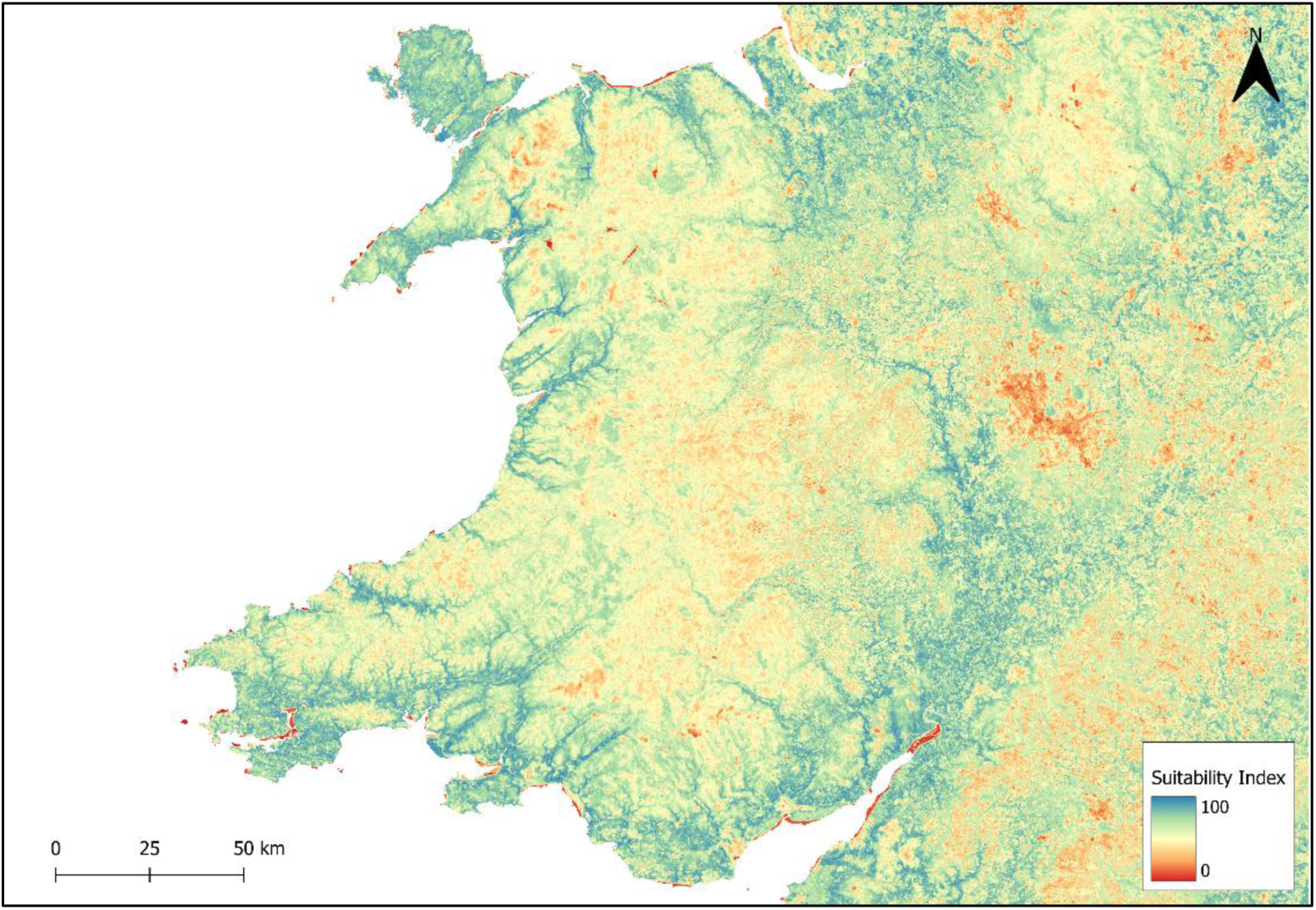

<u>South West England</u>

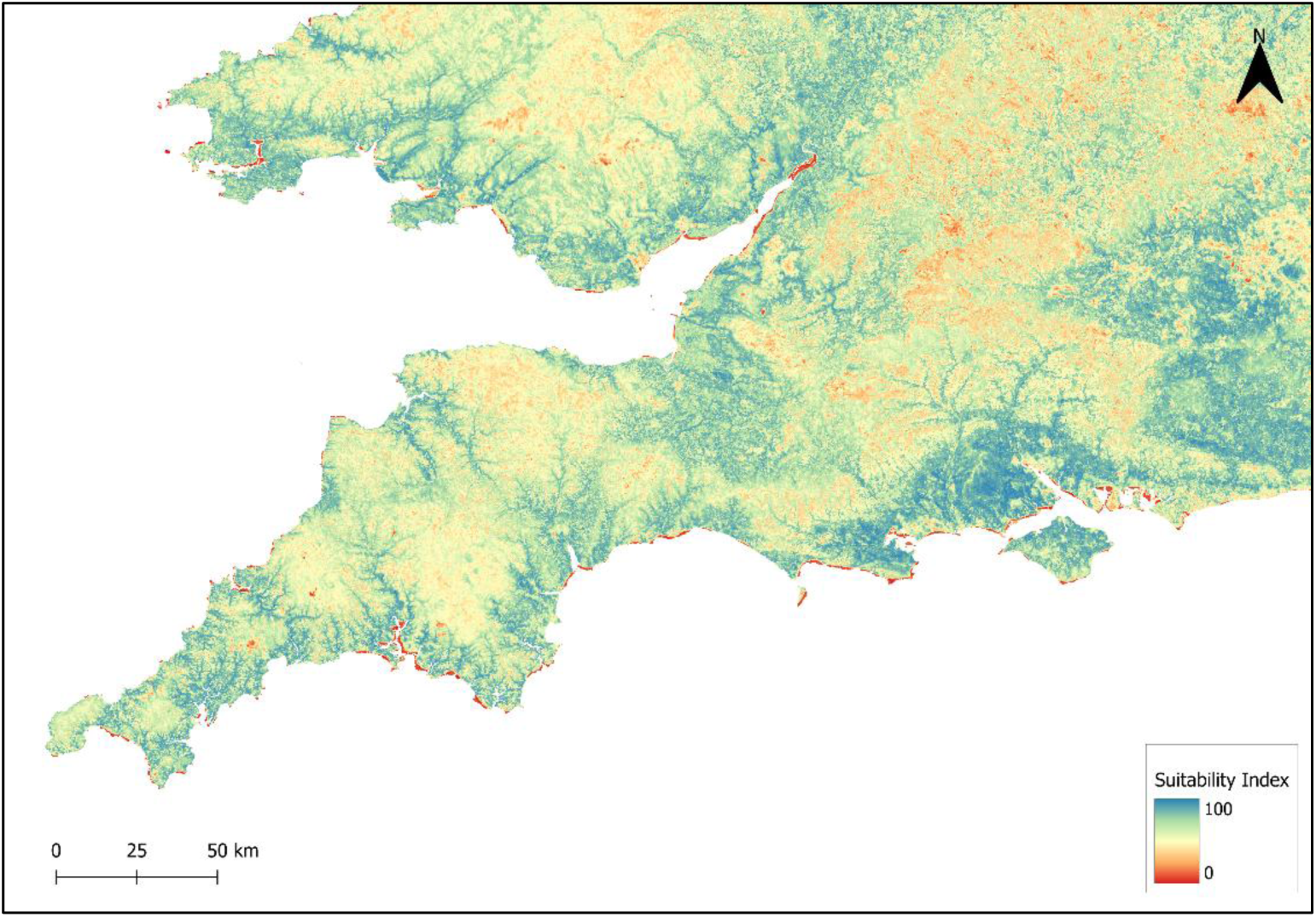

<u>South East England</u>

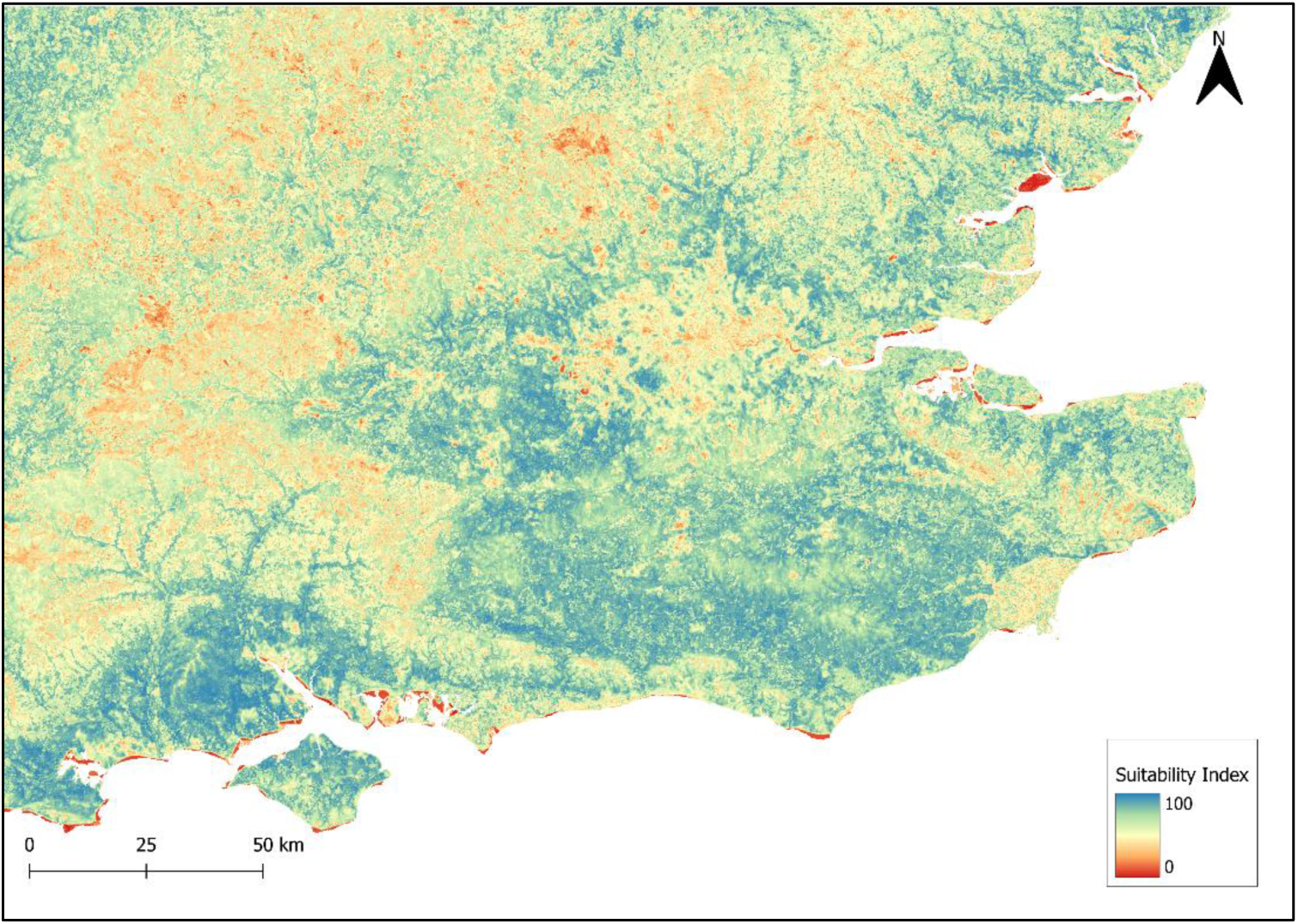

<u>East Anglia</u>

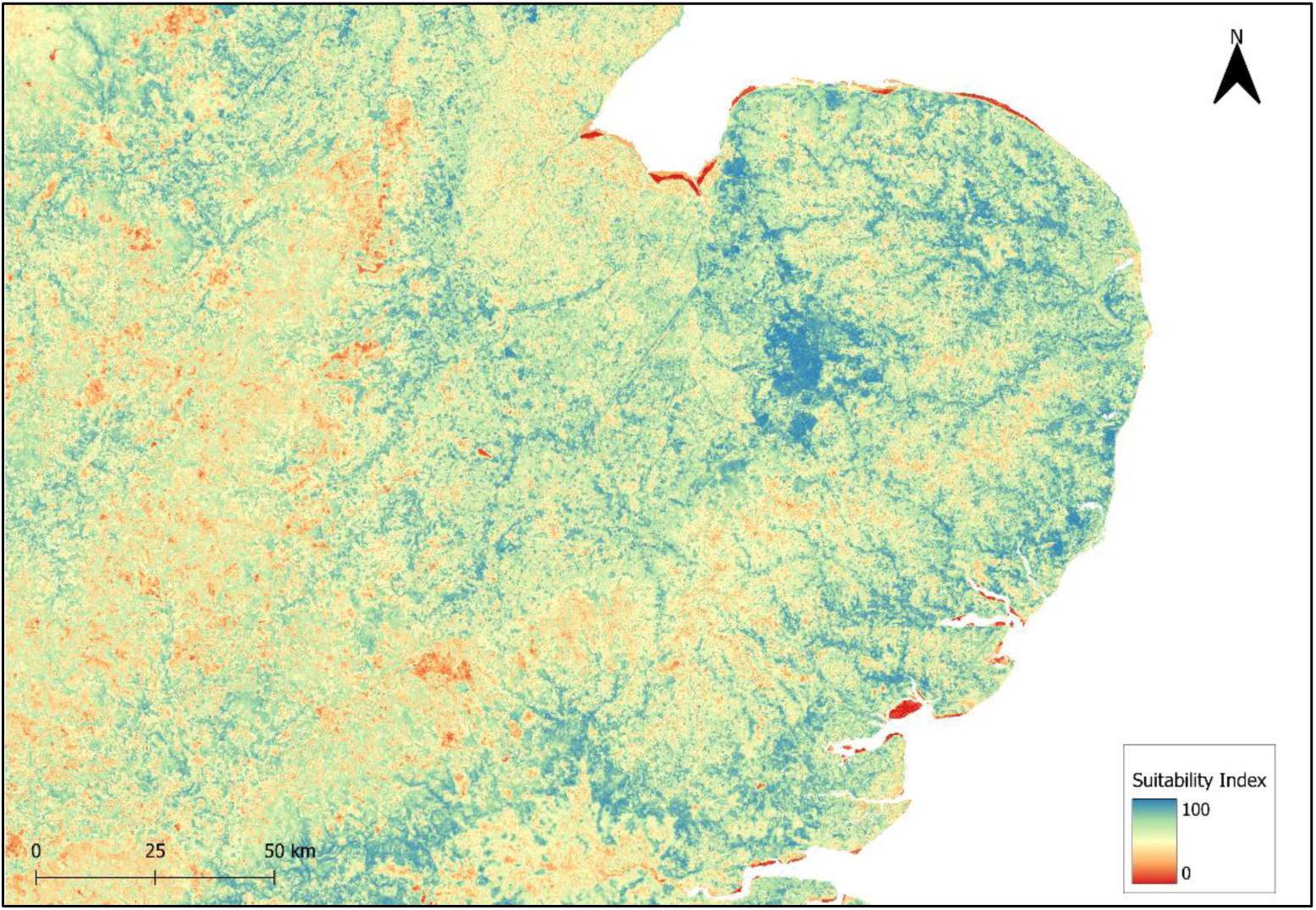

## Appendix 2

Reclassification of CORINE land classes and UKCEH land classes to form 17 land classifications. NA indicates the categories which were not available in the UKCEH layer.

| UKCEH land class number | Alternative layer for UKCEH land class | CORINE land class number | Merged land class categories (combined CORINE and UKCEH) | Classification code of combined categories |
| --- | --- | --- | --- | --- |
| 20 |  | 1 to 9, 11 | Urban | 1 |
| NA | OS Open Greenspace (Ordnance Survey, 2022) | 10 | Urban green | 2 |
| 3 |  | 12, 19 to 22 | Arable | 3 |
| NA |  | 18 | Pastures | 4 |
| 1 |  | 23 | Broadleaf | 5 |
| 2 |  | 24 | Conifer | 6 |
| NA |  | 25 | Mixed woodland | 7 |
| 4 to 7 |  | 26 | Natural grassland | 8 |
| 9 |  | 27 | Moor and heath | 9 |
| NA |  | 28 | Sclerophyllous vegetation | 10 |
| NA |  | 29 | Transitional woodland-scrub | 11 |
| 12 |  | 30 to 34 | Bare (bare rock/sparse veg/burnt area/glacial/perpetual snow) | 12 |
| 8 |  | 35 | Inland marsh | 13 |
| 11 |  | 36 | Peat bogs | 14 |
| 19 |  | 37 | Salt marsh | 15 |
| 13, 15, 16 |  | 38 to 39, 42 to 44 | Coastal | 16 |
| 14 |  | 40 to 41 | Freshwater | 17 |

## Appendix 3

Ensemble model evaluation results from the ensemble model outputs, indicating model performance across TSS, ROC and Kappa evaluations.

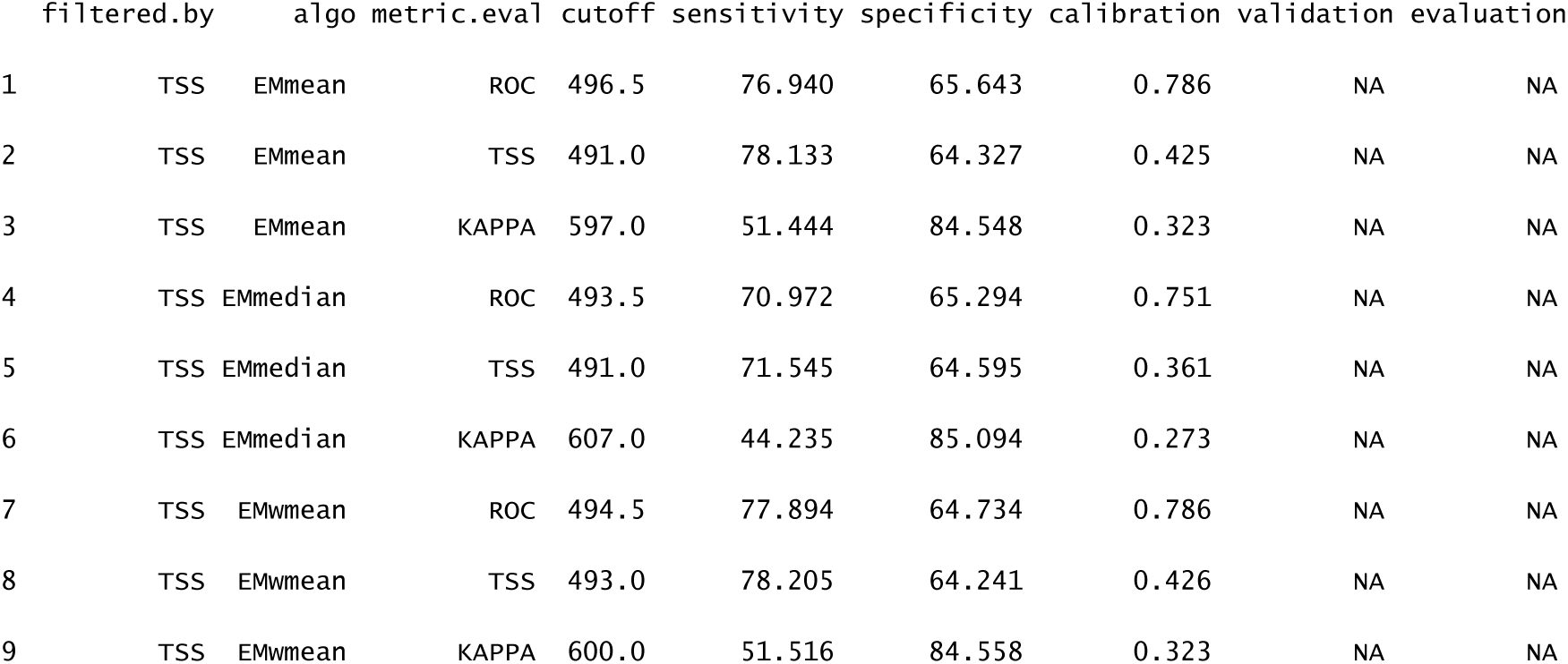

## Appendix 4

Ensemble variable importance results from ensemble model outputs across mean, median and weighted average evaluations.

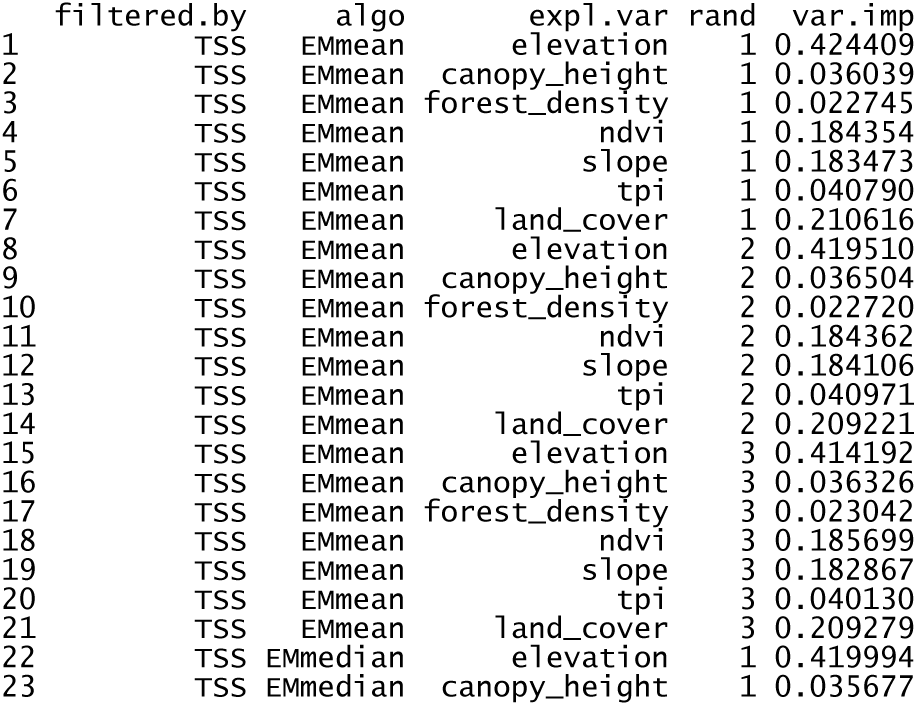
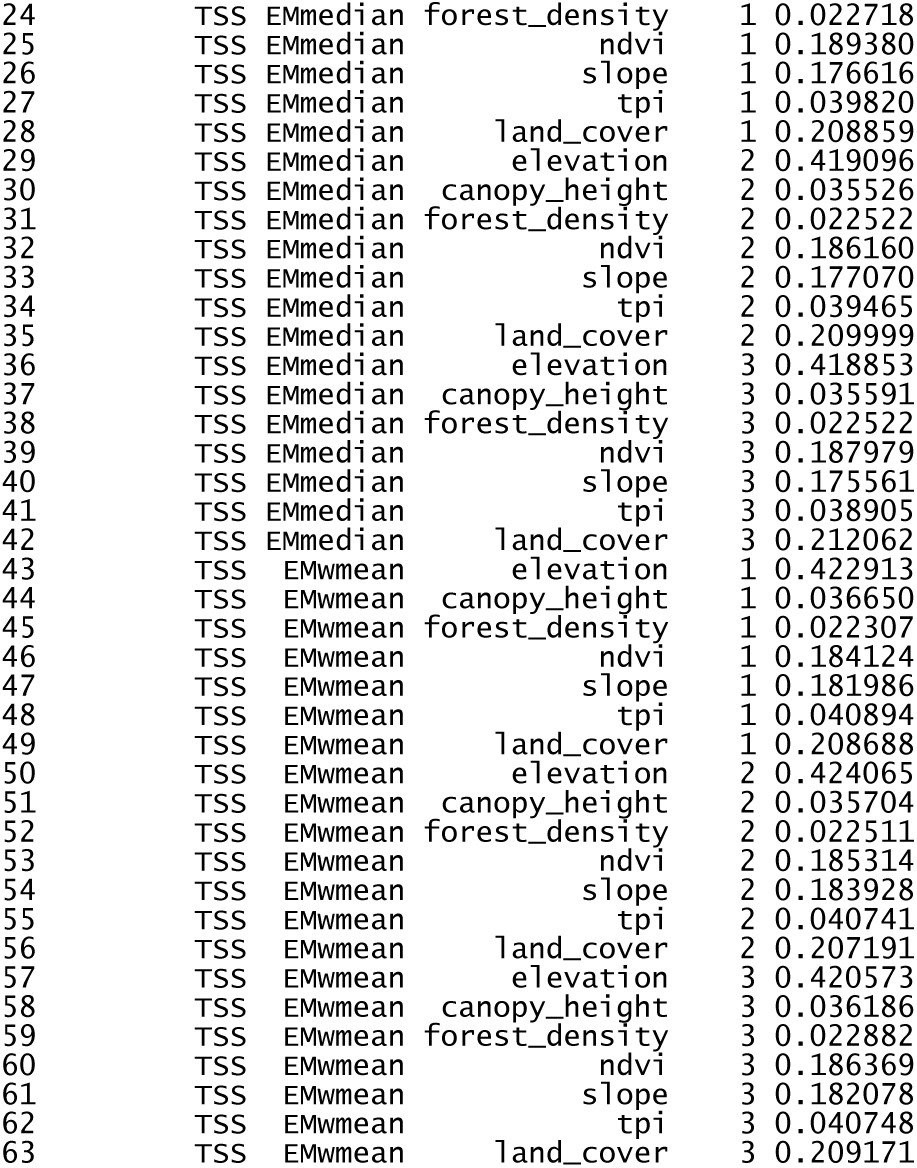

