## Supplementary material for "Is Britain still Great for Pine marten? A Habitat Suitability Assessment": R Script

# Load packages -----------------------

library(sf)  #*
library(dismo)  #*
library(dplyr)  #*
library(raster)  #*
library(tidyr)  #*
library(vegan) #*
library(rgbif)  #*
library(spThin) #*
library(leaflet)
library(geoR)
library(rasterVis)
library(CoordinateCleaner)
library(reshape2)
library(PerformanceAnalytics) 
library(pals)
library(ggplot2)
library(ggpubr)
library(devtools) 
library(tidyterra)
library(ggtext)
library(terra)
library(doParallel)
library(biomod2)

# Create color palettes ------------------------

CB7  <- c("#E69F00", "#56B4E9", "#009E73", "#F0E442", "#0072B2", "#D55E00", "#CC79A7")

CB15 <- c("#000000","#004949","#009292","#ff6db6","#ffb6db",
          "#490092","#006ddb","#b66dff","#6db6ff","#b6dbff",
          "#920000","#924900","#db6d00","#24ff24","#ffff6d")


# Fetch occurrence records from GBIF --------------------
run_code <- FALSE

if(run_code) {
  
  # GBIF credentials stored in r environ
  user <- Sys.getenv("gbif_user")
  pwd <- Sys.getenv("gbif_password")
  email <- Sys.getenv("work_email")
  
  # Define species and run data search
  myspecies <- c("Martes martes")
  
  ?occ_data
  gbif_data <- occ_data(scientificName = myspecies,
                        hasCoordinate = TRUE,
                        hasGeospatialIssue = FALSE,
                        year = '1945,2024',
                        limit=100000,
                        basisOfRecord = "HUMAN_OBSERVATION;
                                        OBSERVATION;
                                        MACHINE_OBSERVATION;
                                        MATERIAL_CITATION;
                                        MATERIAL_SAMPLE;
                                        PRESERVED_SPECIMEN;
                                        OCCURRENCE",
                        coordinateUncertaintyInMeters = '0,1000')
  
  names(gbif_data)
  names(gbif_data$meta)
  names(gbif_data$data)
  
  print(gbif_data$data)
  
  summary(gbif_data$data$coordinateUncertaintyInMeters)
  
  # Save GBIF data as R object.
  saveRDS(gbif_data, file = "Pine Marten/Occurrence data/GBIF_download.rds")
  
  # Restore the object
  #gbif_data1 <- readRDS(file ="Pine Marten/Occurrence data/GBIF_download.rds")
  
  # Subset data part of object
  gbif_data2 <- gbif_data1$data
  head(gbif_data2)
  
  # Look at each column and unlist any lists
  gbif_data3 <- apply(gbif_data2, 2, function(y) sapply(y, function(x) paste(unlist(x), collapse=";")))
  
  # Convert to df
  gbif_data4 <- as.data.frame(gbif_data3)

  # Convert to sf, specify coordinate columns and set crs.
  gbif_data5 <- st_as_sf(gbif_data4, coords = c("decimalLongitude", "decimalLatitude"), crs = 4326, remove=F)

  # Double check CRS is correct
  crs(gbif_data5)
  
  # View coordinates from point geometry
  st_geometry(gbif_data5)
  
  # View coordinates from X and Y columns
  head(gbif_data5$decimalLongitude)
  head(gbif_data5$decimalLatitude)
  
  # Export as geopackage.
  st_write(gbif_data5, "Pine Marten/Occurrence data/GBIF_data.gpkg")
  
  #gbif_data6 <- st_read("Pine Marten/Occurrence data/GBIF_data.gpkg")
  
  gbif_data6$decimalLatitude <- as.numeric(gbif_data6$decimalLatitude)
  gbif_data6$decimalLongitude <- as.numeric(gbif_data6$decimalLongitude)
  gbif_data6$coordinateUncertaintyInMeters <- as.numeric(gbif_data6$coordinateUncertaintyInMeters)
  gbif_data6$eventDate <- as.Date(gbif_data6$eventDate)
  
}

# Fetch occurrence records from NBN Atlas ------------
run_code <- FALSE

if(run_code) {
  
  # Update this url for the species of interest
  search <- "https://records-ws.nbnatlas.org/occurrences/search?q=*:*&fq=genus:Martes&pageSize=20000"
  
  nbn_download <- jsonlite::fromJSON(search, simplifyDataFrame = TRUE) ## the data is pulled into an object called `occurrences`
  
  nbn_data <- nbn_download$occurrences
  
  # Look at each column and unlist any lists
  nbn_data1 <- apply(nbn_data, 2, function(y) sapply(y, function(x) paste(unlist(x), collapse=";")))
  
  # Convert to df
  nbn_data2 <- as.data.frame(nbn_data1)
  
  nbn_data2$eventDate <- as.Date(paste(nbn_data2$year, nbn_data2$month, "01", sep = "-"))
  nbn_data2$coordinateUncertaintyInMeters <- as.numeric(nbn_data2$coordinateUncertaintyInMeters)
  
  # Convert to spatial object, specify coordinate columns, set crs and remove rows with NAs for lat or long
  nbn_data2$decimalLatitude <- as.numeric(nbn_data2$decimalLatitude)
  nbn_data2$decimalLongitude <- as.numeric(nbn_data2$decimalLongitude)
  summary(nbn_data2$decimalLatitude)
  summary(nbn_data2$decimalLongitude)
  nbn_data3 <- subset(nbn_data2, !is.na(decimalLongitude))
  
  # Convert to sf
  nbn_data4 <- st_as_sf(nbn_data3, coords = c("decimalLongitude", "decimalLatitude"), crs = 4326, remove=F)
  
  # Write to geopackage
  st_write(nbn_data4, "Pine Marten/Occurrence data/NBN_data.gpkg", append=F)
  
  #nbn_data4 <- st_read("Pine Marten/Occurrence data/NBN_data.gpkg")
  
  nbn_data5 <- nbn_data4 %>% select(
    decimalLatitude,
    decimalLongitude,
    species = scientificName,
    basisOfRecord,
    eventDate,
    coordinateUncertaintyInMeters,
    institutionCode = raw_institutionCode)
  
}


# Combine GBIF and NBN Atlas occurrence data ---------------------
run_code <- FALSE

if(run_code){
  
  all_data <- bind_rows(gbif_data6, nbn_data5)
  
}


# Data cleaning -----------------------------
run_code <- FALSE
if(run_code){
  
  #We followed the cleaning pipeline outlined by Zizka et al. (2019) and first filtered the data as downloaded from GBIF (“raw”, hereafter) 
  #using meta-data for those records for which they were available (although meta-data were often missing, Peterson et al., 2018), 
  #removing: 
  #(1) records with a coordinate precision below 100 km (as this represents the grain size of many macro-ecological analyses); 
  #(2) fossil records and records of unknown source; 
  #(3) records collected before 1945 (before the end of the Second World War, since coordinates of old records are often imprecise); 
  #and (4) records with an individual count of less than one and more than 99. 
  #Furthermore, we rounded the geographic coordinates to four decimal places and retained only one record per species per location 
  #(i.e., test for duplicated records). In a second step, we used the clean_coordinates function of the CoordinateCleaner v2.0-11 
  #package (Zizka et al., 2019) with default options to flag errors that are common to biological data sets (“filtered”, hereafter). 
  #These include: coordinates in the sea for terrestrial taxa and on land for marine taxa, coordinates containing only zeros, 
  #coordinates assigned to country and province centroids, coordinates within urban areas, and coordinates assigned to biodiversity 
  #institutions. See Table 2 for a summary of all filters we used and their classification into “erroneous” and “unfit”.
  
  # View issues column of occurrence data
  summary(as.factor(all_data$issues))
  
  # Remove rounded coordinates (original coordinate modified by rounding to 5 decimals)
  # The code cdround represents the GBIF issue COORDINATE_ROUNDED
  all_data0 <- subset(all_data$decimalLongitude, nchar(as.character(all_data$decimalLongitude)) > 5)
  summary(as.numeric(all_data$coordinatePrecision))
  
  # View basis of record
  summary(as.factor(all_data$basisOfRecord))
  
  # Filter by date
  summary(as.Date(all_data$eventDate))
  all_data1 <- subset(all_data0, eventDate > "1945-01-01")
  summary(all_data1$eventDate)
  
  # Run clean_occurrences from the CoordinateCleaner package
  ?clean_coordinates
  all_data2 <- clean_coordinates(as.data.frame(all_data1),
                                 lon = "decimalLongitude",
                                 lat = "decimalLatitude",
                                 species = "species",
                                 value="clean")  #requires internet
  
  # Check and filter by coordinateUncertaintyInMeters
  summary(all_data2$coordinateUncertaintyInMeters)
  all_data3 <- subset(all_data2, coordinateUncertaintyInMeters <= 1000)
  summary(all_data3$coordinateUncertaintyInMeters)
  
  # Convert to sf
  all_data4 <- st_as_sf(all_data3, coords = c("decimalLongitude", "decimalLatitude"), crs = 4326, remove=F)
  
  # Save the cleaned occurrences
  st_write(all_data4, "Pine Marten/Occurrence data/GBIF_NBN_data.gpkg")
  
  # Import the cleaned occurrence data 
  gbif_nbn_data <- st_read("Pine Marten/Occurrence data/GBIF_NBN_data.gpkg")
  
}


# Read in analysis area shapefile ---------------------
europe_boundary <- st_read("Pine Marten/AOI shapefile/europe_shapefile.gpkg")
crs(europe_boundary)
plot(europe_boundary[1])


# Fetch or add environmental variables raster data --------------------
run_code <- FALSE
if(run_code){
  
  # Import and resample, align, crop and mask rasters. Masks everything outside the polygons as nodata. 
  
  setwd("Pine Marten/Environmental rasters/rasters_eu")
  
  # Vector of file names
  tif_files <- list.files(pattern='\\.tif$')
  tif_files
  tif_files <- tif_files[c(1,3,4,5,6,8,10)]
  tif_files
  
  # Import rasters
  myrasters <- lapply(tif_files, raster) #requires internet
  myrasters
  
  # Plot a raster
  plot(myrasters[[3]])
  
  # Subset the rasters to discrete and continuous
  myrasterscontinuous <- myrasters[c(1,3,5,6,7)] # exclude reference raster here
  myrastersdiscrete <- myrasters[c(4)]
  myrasterscontinuous
  myrastersdiscrete
  
  
  # Subset the raster to be used as the reference
  myrasters
  ref_raster <- myrasters[[2]]
  ref_raster
  
  # Set discrete rasters to factor
  myrastersdiscrete1 <- lapply(myrastersdiscrete, as.factor)
  myrastersdiscrete
  myrastersdiscrete1
  
  # Resample other rasters to match reference raster and align extent.
  myrasterscontinuous_resampled <- lapply(myrasterscontinuous, function(x){raster::resample(x, ref_raster, method="bilinear")})
  myrastersdiscrete_resampled <- lapply(myrastersdiscrete1, function(x){raster::resample(x, ref_raster, method="ngb")})
  
  # Set discrete rasters to factor
  myrastersdiscrete_resampled1 <- lapply(myrastersdiscrete_resampled, as.factor)
  
  # Combine the rasters
  myrasters1 <- c(ref_raster, myrasterscontinuous_resampled, myrastersdiscrete_resampled)
  myrasters1
  plot(myrasters1[[1]])
  plot(europe_boundary, add=T, col=NA)
  
  # Crop and mask rasters to AOI shapefile
  myrasters2 <- lapply(myrasters1, function(x){crop(x, europe_boundary)})
  myrasters3 <- lapply(myrasters2, function(x){mask(x, europe_boundary)})
  
  # Plot a raster
  plot(myrasters3[[7]])
  
  # Stack rasters
  mystack <- stack(myrasters3)
  mystack@layers
  
  # Set discrete rasters to factor
  mystack[[7]]
  mystack[[7]] <- as.factor(mystack[[7]])
  mystack@layers
  
  # Save the resampled stacked rasters as a grd file
  setwd("Pine Marten/Environmental rasters")
  mystack_pro <- mystack
  writeRaster(mystack_pro, "rasterstack_eu", format="raster", overwrite=TRUE)
  
}

# Import the Europe raster stack ------------------------
run_code <- TRUE
if(run_code){
  
  setwd("Pine Marten/Environmental rasters")
  
  mystack_pro <- stack("rasterstack_eu.grd")
  mystack_pro@layers
  
  # Set discrete rasters to factor
  mystack_pro[[7]]
  mystack_pro[[7]] <- as.factor(mystack_pro[[7]])
  mystack_pro@layers
  
}

# Remove occurrences outside AOI, check collinearity, and spatial thinning ----------------
run_code <- FALSE

if(run_code){
  
  # Reproject the occurrence data to match the raster data
  spdata <- gbif_nbn_data %>% st_transform(crs(mystack_pro))
  crs(spdata)
  crs(mystack_pro)
  
  # Extract environmental variables from raster to points ----------------------
  envtopoint <- raster::extract(mystack_pro, spdata, df=T)
  spdata <- spdata %>% rename(elevation_gbifnbn = "elevation")
  spdata1 <- cbind(spdata, envtopoint)
  
  # Identify the environmental variables columns
  colnames(spdata1)
  spdata1[169:175]
  
  # Remove points that lie outside of the analysis area.
  spdata2 <- drop_na(spdata1, c(169:175))
  
  # Remove duplicates based on lat, long and species data
  # Check species column values
  unique(spdata2$species)
  # Update if needed
  spdata2$species <- "Martes martes"
  summary(spdata2$decimalLongitude)
  summary(spdata2$decimalLatitude)
  # Remove duplicates
  ?distinct
  spdata3 <- spdata2 %>% dplyr::distinct(decimalLongitude, decimalLatitude, .keep_all=T)
  
  # Plot histograms of environmental variables
  # Specify the range of variables (columns 169 to 175)
  vars_to_plot <- st_drop_geometry(spdata3[, 169:175])
  # Loop through each column and create histograms
  for (i in seq_along(vars_to_plot)) {
    # Get the column name to use in the file name
    var_name <- names(vars_to_plot)[i]
    # Create a PDF file for each histogram
    pdf(file = paste0("Pine Marten/Other outputs/", var_name, "_histogram.pdf"))
    # Create the histogram
    hist(vars_to_plot[[i]], 
         main = paste("Histogram of", var_name), 
         xlab = var_name, 
         col = "lightblue", 
         border = "black")
    # Close the PDF file
    dev.off()
  }
  
  # Summary of distances between points (m) for a given species
  head(st_drop_geometry(spdata3[,c("decimalLongitude","decimalLatitude")]))
  head(st_coordinates(spdata3))
  spdata3$X3035 <- st_coordinates(spdata3)[, 1]
  spdata3$Y3035 <- st_coordinates(spdata3)[, 2]
  ?dist
  dists <- dist(st_coordinates(spdata3))
  summary(dists/1000)
  
  # Check for collinearity in a correlation matrix
  spdata3a <- st_drop_geometry(spdata3)
  names(spdata3a)
  summary(spdata3a[169:175]) # check for NAs - shouldnt be any as removed with drop_na earlier.
  chart.Correlation(spdata3a[169:175], histogram=TRUE, pch=19)
  
  # Create geoR geodata object for variogram. 
  # This will remove points which have NA in some of the rasters, for example at the coastline
  summary(spdata3a[176:177])
  
  datageo <- as.geodata(spdata3a,
                        coords.col = 176:177,
                        data.col = 169:174)  # cannot computer variogram on factor variables, so leave them out.
  summary(datageo)
  
  # Set lags for variogram. We want >= 30 pairs per lag (v1$n). 
  # Adjust the sequence and l value until you get a sensible plot output. 
  #breaks = seq(0, 100, l = 50) #also tried (0,500, 1 = 250)
  #breaks
  
  # Create variogram. 
  # Set the max distance of interest. 
  ?variog
  v1 <- variog(datageo,
               max.dist = 30000)
  summary(v1)
  v1$n
  v1$u
  v1$u <- v1$u/1000
  v1$u
  
  # Plot variogram
  
  png(filename = "Pine Marten/Other outputs/Variogram.png",width = 1600, height = 1200, units = "px", pointsize = 30)
  par(mar=c(5,5,5,9))
  par(xpd=T)
  
  plot(v1, type="l", scaled=T, lwd=2, pch=20, cex=0.2, lty=1, col=CB7,
       ylim=c(0,1),
       xlim=c(0,30),
       xlab= 'Distance (km)',
       ylab = 'Semivariance',
  )
  legend("topright", legend=levels(as.factor(names(v1$var.mark))), lty=1, col=CB7,
         cex=0.7,
         #inset=c(-0.2,0),
         lwd=2)
  par(xpd=F)
  abline(v=c(10), col=c("black"), lty=c(2,2), lwd=c(3, 3))
  text(x=15, y=0.1, labels="practical range: 10 km")
  dev.off()
  
  # Perform thinning, which returns only coordinates
  setwd("Pine Marten/Occurrence data/SpThin")
  unique(spdata3a$species)
  
  # Set seed
  set.seed(1)
  ?thin
  spdata3a_thin <- thin(spdata3a,
                        lat.col = "decimalLatitude",
                        long.col = "decimalLongitude",
                        spec.col = "species",
                        thin.par = 10, # decide on this based on variogram etc. This is in kilometres!
                        reps = 30,
                        locs.thinned.list.return = TRUE, #gives the list of dataframes for each repetition iteration, so 30 dataframes
                        write.files = T,
                        max.files = 1,
                        out.dir = "Pine Marten/Occurrence data/SpThin",
                        out.base = "thinned_data",
                        write.log.file = T,
                        log.file = "Pine Marten/Occurrence data/SpThin/thinout.txt",
                        verbose = TRUE)
  
  # Select the first thinning repetition which has retained the most occurrence data. 
  spdata3a_thin[1]
  spdata3b <- as.data.frame(spdata3a_thin[1])
  head(spdata3b)
  
  # Merge the thinned occurrences data with the original columns.
  ?merge
  spdata4 <- merge(spdata3, spdata3b, by="row.names")
  spdata4
  crs(spdata4)
  
  # Plot the thinned data
  mystack_pro[[4]]
  plot(mystack_pro[[4]])
  spdata4[175]
  plot(spdata4[175], pch=20, cex=1, add=T)
  
  # Export and import thinned occurrence data
  setwd("Pine Marten/Occurrence data")
  saveRDS(spdata4, file = "GBIF_NBN_data_thinned.rds")
  
}

# Import the cleaned and thinned occurrence data -------------------
run_code <- TRUE
if(run_code){
  
  setwd("Pine Marten/Occurrence data")
  spdata4 <- readRDS(file = "GBIF_NBN_data_thinned.rds")

}

# Prepare data for BIOMOD2 Species Distribution Modelling (SDM) -----------------------
run_code <- TRUE
if(run_code){
  
  # Number of occurrences after cleaning and thinning
  nrow(spdata4)
  
  # Reformat occurrences to format required by BIOMOD2
  spdata5 <- spdata4
  spdata5$constant <- rep(1,nrow(spdata5))
  spdata5$constant
  
  # Set WD for modelling
  setwd("Pine Marten/Biomod")
  
  # Define BIOMOD2 inputs to match vignette terminology
  myRespName <- 'Martes1'
  myResp <- st_drop_geometry(spdata5$constant)
  st_drop_geometry(spdata5[c('X3035','Y3035')])
  myRespCoord = st_drop_geometry(spdata5[c('X3035','Y3035')]) 
  mystack_pro@layers
  myExpl <- mystack_pro
  myExpl@layers
  
  # Choose number of pseudoabsences (PA)
  no.pa <- sum(myResp == 1, na.rm=T) # Same number as presences
  no.pa
  
}


# Load in previously produced formal models --------------------
run_code <- FALSE
if(run_code){
  
  setwd("Pine Marten/Biomod")
  list.files("Martes1", recursive = T)
  
  bm_out_file <- load("Martes1/Martes1.1729527507.models.out")
  myBiomodModelOut <- get(bm_out_file)
}

# Load in previously produced ensemble models --------------------
run_code <- FALSE
if(run_code){
  
  setwd("Pine Marten/Biomod")
  list.files("Martes1", recursive = T)
  
  bm_ensemble_out_file <- load("Martes1/Martes1.1729527507.ensemble.models.out")
  myBiomodEM <- get(bm_ensemble_out_file)
}

# Load in previously produced formal model projections for Europe --------------------
run_code <- FALSE
if(run_code){
  
  setwd("Pine Marten/Biomod")
  list.files("Martes1", recursive = T)
  
  bm_proj_out_file <- load("Martes1/proj_em_uk_lb/Martes1.em_uk_lb.ensemble.projection.out") 
  myBiomodProj_EU <- get(bm_proj_out_file)
}

# Load in previously produced ensemble model projections for Europe --------------------
run_code <- FALSE
if(run_code){
  
  setwd("Pine Marten/Biomod")
  list.files("Martes1", recursive = T)
  
  bm_em_proj_out_file <- load("Martesmartes/proj_current_ensemble3RUNS/Martesmartes.current_ensemble3RUNS.ensemble.projection.out")
  myBiomodEF <- get(bm_em_proj_out_file)
}

# Load in previously produced formal model projections for Britain --------------------
run_code <- FALSE
if(run_code){
  
  setwd("Pine Marten/Biomod")
  list.files("Martes1", recursive = T)
  
  bm_proj_out_file <- load("Martesmartes/proj_100m_Britain_3RUNS/Martesmartes.100m_Britain_3RUNS.projection.out") #won't work
  myBiomodProj_UK <- get(bm_proj_out_file)
}

# Load in previously produced ensemble model projections for Britain --------------------
run_code <- FALSE
if(run_code){
  
  setwd("Pine Marten/Biomod")
  list.files("Martes1", recursive = T)
  
  bm_em_proj_out_file <- load("Martesmartes/proj_current_el_Britain/Martesmartes.current_el.ensemble.projection.out")
  myBiomodEFBritain <- get(bm_em_proj_out_file)
}


# Formatting data - BIOMOD2 Species Distribution Modelling (SDM) -----------------------
run_code <- TRUE
if(run_code){
  
  ?BIOMOD_FormatingData
  myBiomodData <- BIOMOD_FormatingData(resp.name = myRespName, # the species name
                                       resp.var = myResp, # the binary presence/absence occurrence data 
                                       expl.var = myExpl, # the raster stack
                                       resp.xy = myRespCoord, # the occurrence data coordinates
                                       eval.resp.var = NULL, # the binary presence/absence independent testing data
                                       eval.resp.xy = NULL, # the independent testing occurrence data coordinates
                                       eval.expl.var = NULL, # the independent testing raster data
                                       PA.nb.rep = 2, # PA repetitions created. 
                                       PA.nb.absences = no.pa, # number of PAs (currently, same as the number of presences)
                                       PA.strategy = 'disk', # strategy to select the PAs
                                       PA.dist.min = 5000, # minimum distance to presence points for disk PA selection
                                       PA.dist.max = NULL, # maximum distance to presence points for disk PA selection
                                       na.rm = T # whether points having any missing values for explanatory variables should be removed
  )
  
  plot(myBiomodData)
  show(myBiomodData)
}


# Define options - BIOMOD2 Species Distribution Modelling (SDM) -----------------------
run_code <- TRUE
if(run_code){
  
  # Currently using the default options
  ?BIOMOD_ModelingOptions
  myBiomodOptions <- BIOMOD_ModelingOptions() # default model settings. E.g. glm is default to quadratic, AIC and binomial family 
}

########################################################
# cl <- makeCluster(detectCores() - 1)  # Use one less than the number of available cores
# registerDoParallel(cl)
# stopCluster(cl)
#######################################################

# Build formal models - BIOMOD2 Species Distribution Modelling (SDM) -----------------------
run_code <- TRUE
if(run_code){
  
  ?BIOMOD_Modeling
  myBiomodModelOut <- BIOMOD_Modeling(
    bm.format = myBiomodData, 
    models = c('GLM',   
               'GAM',
               'GBM',
               'CTA',
               'ANN',
               'SRE',
               'FDA',
               'MARS',
               'RF', 
               'MAXNET',
               "XGBOOST"
    ), 
    models.pa = NULL, # a list of pseudo-absence datasets for each model
    bm.options = myBiomodOptions,
    CV.strategy = "random", # the cross-validation selection strategy
    CV.nb.rep = 2, # the number of sets (repetitions) of cross-validation points that will be drawn
    CV.perc = 0.7, # the test-training data percentage split
    CV.k = NULL, # an integer corresponding to the number of partitions (kfold, strat or env)
    CV.balance = NULL, # how data will be balanced between partitions (strat or env)
    CV.env.var = NULL, # the environmental variables used to build the partition (env)
    CV.strat = NULL, # how data will be partitioned along gradient (env)
    CV.user.table = NULL, # defining for each repetition (in columns) which observation lines should be used for models calibration (TRUE) and validation (FALSE) (user.defined)
    CV.do.full.models = FALSE, # whether models should be also calibrated and validated over the whole dataset (and pseudo-absence datasets)
    weights = NULL,  # null means that presences and PA's are weighted equally, which is what we want according to literature
    prevalence=0.5, #species prevalence to build 'weighted response weights', standard is 0.5
    metric.eval = c('ROC', 'TSS', 'KAPPA'), # evaluation metrics to be used: ROC, TSS, KAPPA, ACCURACY, BIAS, POD, FAR, POFD, SR, CSI, ETS, HK, HSS, OR, ORSS
    var.import=3, # number of permutations to estimate variable importance, has been either 0 or 3 from examples, a table which says how important each raster is to predict habitat suitability (predictive power).
    scale.models = FALSE, # whether all models predictions should be scaled with a binomial GLM or not
    nb.cpu = 24,
    seed.val = 2,
    do.progress = TRUE)
  
  show(myBiomodModelOut)
  
  # Save model evaluation scores and variables importance on hard drive
  capture.output(get_evaluations(myBiomodModelOut), 
                 file=file.path(paste(myRespName,"_formal_models_evaluation.txt", sep="",options(max.print = 5000))))
  capture.output(get_variables_importance(myBiomodModelOut), 
                 file=file.path(paste(myRespName,"_formal_models_variable_importance.txt", sep="")))
}
  

# View formal models evaluation and variable importance -------------------
run_code <- TRUE
if(run_code){
  variable_importance <- get_variables_importance(myBiomodModelOut) 
  variable_importance
  
  model_scores <- get_evaluations(myBiomodModelOut)
  model_scores
  
  summary(subset(model_scores, metric.eval == "KAPPA")$validation)

  quantile(subset(model_scores, metric.eval == "TSS")$validation)
  unname(quantile(subset(model_scores, metric.eval == "TSS")$validation)[4])
  
  ?bm_PlotEvalMean # mean evaluation scores (and their standard deviation)
  bm_PlotEvalMean(bm.out = myBiomodModelOut)
  
  bm_PlotEvalBoxplot
  bm_PlotEvalBoxplot(bm.out = myBiomodModelOut, dataset = "validation", group.by = c('algo', 'algo'))
  bm_PlotEvalBoxplot(bm.out = myBiomodModelOut, group.by = c('algo', 'run'))
  
  ?bm_PlotVarImpBoxplot
  bm_PlotVarImpBoxplot(bm.out = myBiomodModelOut, group.by = c('expl.var', 'algo', 'algo'))
  bm_PlotVarImpBoxplot(bm.out = myBiomodModelOut, group.by = c('expl.var', 'algo', 'run'))
  bm_PlotVarImpBoxplot(bm.out = myBiomodModelOut, group.by = c('expl.var', 'algo', 'PA'))
  
}


# Build ensemble models - BIOMOD2 Species Distribution Modelling (SDM) -----------------------
run_code <- TRUE
if(run_code){
  
  ?BIOMOD_EnsembleModeling
  myBiomodEM <- BIOMOD_EnsembleModeling(
    bm.mod = myBiomodModelOut, 
    models.chosen = 'all', # a vector containing model names to be kept
    em.by= 'all',  # the way in which the models will be combined, the default is 'PA+run', 'all' means it includes all the data before the splitting. 
    em.algo = c('EMmean', 'EMmedian', 'EMwmean'), 
    metric.select = 'TSS', # metric names to be used together with metric.select.thresh to exclude single models based on their evaluation scores
    metric.select.thresh = unname(quantile(subset(get_evaluations(myBiomodModelOut), metric.eval == "TSS")$validation)[4]),  # A vector of numeric values corresponding to the minimum scores (one for each metric.select) below which single models will be excluded from the ensemble model building 
    metric.select.table = NULL, # evaluation scores calculated for each single models and that will be compared to metric.select.thresh values to exclude some of them from the ensemble model building
    metric.select.dataset = NULL, # which dataset should be used to filter and/or weigh the ensemble models should be among 'evaluation', 'validation' or 'calibration'.
    metric.eval = c('ROC', 'TSS', 'KAPPA'), # evaluation metrics to be used
    var.import = 3, # number of permutations to estimate variable importance
    EMci.alpha = 0.05, # the significance level to estimate confidence interval
    EMwmean.decay = "proportional", # the relative importance of the weights (if 'EMwmean' was given to argument em.algo). A high value will strongly discriminate good models from the bad ones
    nb.cpu = 16,
    seed.val = 3,
    do.progress = TRUE)
  
  
  # Save model evaluation scores and variables importance on hard drive
  capture.output(get_evaluations(myBiomodEM), 
                 file=file.path(paste(myRespName,"_formal_models_ENSEMBLE_evaluation.txt", sep="")))
  capture.output(get_variables_importance(myBiomodEM), 
                 file=file.path(paste(myRespName,"_formal_models_variable_importance_ENSEMBLE.txt", sep="")))
}

# View ensemble model evaluation and variable importance
run_code <- TRUE
if(run_code){
  
 
  
  variable_importance <- get_variables_importance(myBiomodEM) 
  variable_importance
  
  png(filename = "Pine Marten/Other outputs/variable_importance.png", width=2000, height=1000, units = "px", res=300)
  
  ggplot(variable_importance) + 
    geom_boxplot(aes(x=expl.var, y=var.imp)) + 
    labs(x = "Explanatory variable", y = "Importance") + 
    scale_x_discrete(labels = c('Canopy height',
                                'Elevation',
                                'Forest density',
                                'Land cover',
                                'NDVI',
                                'Slope',
                                'TPI')) + 
    theme_minimal()
  
  dev.off()
  
  
  
  model_scores <- get_evaluations(myBiomodEM)
  model_scores
  
  ?bm_PlotEvalMean # mean evaluation scores (and their standard deviation)
  bm_PlotEvalMean(bm.out = myBiomodEM) 
  
  ?bm_PlotEvalBoxplot
  bm_PlotEvalBoxplot(bm.out = myBiomodEM, group.by = c('algo', 'algo'))
  bm_PlotEvalBoxplot(bm.out = myBiomodEM, group.by = c('algo', 'merged.by.run'))
  bm_PlotEvalBoxplot(bm.out = myBiomodEM, group.by = c('algo', 'merged.by.PA'))
  bm_PlotEvalBoxplot(bm.out = myBiomodEM, group.by = c('merged.by.run', 'merged.by.PA'))
  
  ?bm_PlotVarImpBoxplot
  bm_PlotVarImpBoxplot(bm.out = myBiomodEM, group.by = c('expl.var', 'algo', 'merged.by.PA'))
  bm_PlotVarImpBoxplot(bm.out = myBiomodEM, group.by = c('expl.var', 'expl.var', 'algo'))
  
  # Check the TSS score and how many models were kept for ensemble
  get_kept_models(myBiomodEM)
  summary(as.factor(subset(get_evaluations(myBiomodModelOut), metric.eval=="TSS")[10] > 0.324))
  summary(subset(get_evaluations(myBiomodModelOut), metric.eval=="TSS")[10])
  
}

# Import the Britain rasters ----------------------------
run_code <- FALSE
if(run_code){
  
  setwd("Pine Marten/AOI shapefile")
  uk_boundary <- st_read("uk_shapefile_27700.shp") %>% st_transform(3035)
  crs(uk_boundary)
  plot(uk_boundary[1])
  
  setwd("Pine Marten/Environmental rasters/rasters_uk")
  
  # Vector of file names
  tif_files_uk <- list.files(pattern='\\.tif$')
  tif_files_uk
  tif_files_uk <- tif_files_uk[c(1,2,3,4,5,7,9)]
  tif_files_uk
  
  # Import rasters
  myrasters_uk <- lapply(tif_files_uk, raster)
  myrasters_uk
  
  # Subset the raster to be used as the reference
  ref_raster_uk <- myrasters_uk[[2]]
  ref_raster_uk
  
  # Subset the rasters to discrete and continuous
  myrasterscontinuous_uk <- myrasters_uk[c(1,3,5,6,7)]
  myrastersdiscrete_uk <- myrasters_uk[c(4)]
  myrasterscontinuous_uk
  myrastersdiscrete_uk
  
  # Resample other rasters to match reference raster and align extent.
  myrasterscontinuous_uk_resampled <- lapply(myrasterscontinuous_uk, function(x){raster::resample(x, ref_raster_uk, method="bilinear")})
  myrastersdiscrete_uk_resampled <- lapply(myrastersdiscrete_uk, function(x){raster::resample(x, ref_raster_uk, method="ngb")})
  
  # Combine the rasters
  myrasters_uk1 <- c(ref_raster_uk, myrasterscontinuous_uk_resampled, myrastersdiscrete_uk_resampled)
  myrasters_uk1
  
  # Plot boundary over raster
  plot(myrasters_uk1[[1]])
  plot(uk_boundary, add=T, col=NA)
  
  # Crop and mask to AOI polygon 
  myrasters_uk2 <- lapply(myrasters_uk1, function(x){crop(x, uk_boundary)})
  myrasters_uk3 <- lapply(myrasters_uk2, function(x){mask(x, uk_boundary)})
  
  # Stack rasters
  mystack_uk <- stack(myrasters_uk3)
  mystack_uk@layers

  # Compare Europe and UK raster order and names
  names(mystack_pro)
  mystack_pro
  names(mystack_uk)
  mystack_uk
  
  # Set discrete raster as factor
  mystack_uk[[7]] <- as.factor(mystack_uk[[7]])
  levels(mystack_uk[[7]])
  mystack_uk@layers
  mystack_uk[[7]][mystack_uk[[7]] == -9999] <- NA
  mystack_uk@layers

  # Save the resampled stacked rasters as a grd file
  setwd("Pine Marten/Environmental rasters")
  writeRaster(mystack_uk, "rasterstack_uk", format="raster", overwrite=TRUE)
  
}


# Import the Britain raster stack ------------------------
run_code <- TRUE
if(run_code){
  
  setwd("Pine Marten/Environmental rasters")
  
  mystack_uk <- stack("rasterstack_uk.grd")
  mystack_uk@layers
  
  # Set discrete rasters to factor
  mystack_uk[[7]]
  mystack_uk[[7]] <- as.factor(mystack_uk[[7]])
  mystack_uk@layers
  
}

# Projecting formal models ---------------------------
run_code <- TRUE
if(run_code){
  
  setwd("Pine Marten/Biomod")
  
  # Make projection on British 100m rasters
  ?BIOMOD_Projection
  myBiomodProj_UK <- BIOMOD_Projection(
    bm.mod = myBiomodModelOut, 
    proj.name = 'uk_lb', 
    new.env = mystack_uk, # specify here different rasters if projecting to a different area or set of rasters (e.g. climate change study), expl2 is the british rasters
    new.env.xy = NULL,
    models.chosen = 'all',
    metric.binary = 'TSS', #used to be binary.meth
    metric.filter = NULL,
    compress = TRUE, 
    #output.format = ".img",
    build.clamping.mask = TRUE, 
    nb.cpu = 16,
    seed.val = 4)
  
}


# ?get_built_models
#
# mods <- get_built_models(obj = myBiomodModelOut, 
#                          #full.name = NULL, 
#                          PA = 'PA1',
#                          run = 'RUN1')
# mods                    

plot(myBiomodProj_UK, str.grep = 'GLM')
#plot(myBiomodProjEuropeRasters, str.grep = "Martesmartes_PA1_RUN1_GLM")
#plot(myBiomodProjEuropeRasters)

# #cropping europe rasters to britain shapefile for projection
# setwd(wd7)
# tif_files2 <- list.files(pattern='\\.tif$')
# tif_files2
# myrasters <- lapply(tif_files2, raster) 
# myrasters <- myrasters[c(7,1,3,5,6,8,9,10,2,4)]
# myrasters

# setwd(wd6)
# britain_shapefile <-st_read("uk_shapefile.shp") 
# cropped_rasters <- lapply(myrasters, function(x){crop(x, britain_shapefile)})
# cropped_masked_rasters <- lapply(cropped_rasters, function(x){mask(x, britain_shapefile)}) 

# plot(cropped_masked_rasters[[10]])
# cropped_masked_rasters
# europestack1 <- stack(cropped_masked_rasters)
# levels(mystack[[9]])
# levels(mystack[[10]])
# factor(mystack[[9]], level(c(0,1,2,3,4,5,6,7,8,9,10,11,12,13,14,15,16,17,18,19,20))) # assign levels to match European stack, make right way round
# factor(mystack[[9]], level(c(1,2,3,4,5,6,7,8,9,10,11,12,13,14,15,16,17)))
# #europestack1[[9]] <- as.factor(europestack1[[9]])
# europestack1[[9]]
# #europestack1[[10]] <- as.factor(europestack1[[10]])
# europestack1[[10]]

# names(europestack1) <- tif_files3 #make sure the names don't have $ 
# europestack1@layers
# MyExpl3 <- europestack1
# MyExpl3@layers

# Projecting formal models on 1km Europe rasters ---------------------------
run_code <- TRUE
if(run_code){
  
  setwd("Pine Marten/Biomod")
  
  ?BIOMOD_Projection
  myBiomodProj_EU <- BIOMOD_Projection(
    bm.mod = myBiomodModelOut, 
    proj.name = 'eu_proj_lb', 
    new.env = myExpl, # specify here different rasters if projecting to a different area or set of rasters (e.g. climate change study), expl2 is the british rasters
    new.env.xy = NULL,
    models.chosen = "all", 
    metric.binary = 'TSS', #used to be binary.meth
    metric.filter = NULL,
    compress = TRUE, 
    #output.format = ".img", 
    build.clamping.mask = TRUE, 
    nb.cpu = 16,
    seed.val = 1
  )
  
}


# #visualising the projections
# plot(myBiomodProj_EU, str.grep = 'GLM')
# get_built_models(myBiomodModelOut)[22]
# myBiomodProj@proj.out@val@layers
# myBiomodProj@proj.out@val@layers[[2]]
# plot(myBiomodProj@proj.out@val@layers[[1]])


# Projecting ensemble models ---------------------------
run_code <- TRUE
if(run_code){
  
  setwd("Pine Marten/Biomod")
  
  # Make projection on British 100m rasters
  ?BIOMOD_EnsembleForecasting
  myBiomodEF_UK <- BIOMOD_EnsembleForecasting(
    bm.em = myBiomodEM,
    bm.proj = myBiomodProj_UK,
    proj.name = 'em_uk_lb',
    #new.env = myExpl2, #these are the 100m british rasters
    #new.env.xy = NULL,
    models.chosen = 'all', 
    metric.binary = 'TSS', #vector specifying the names of evaluation metrics and associated thresholds to transform the probabilities of presence into presence and absence. Probably TSS. 
    metric.filter = NULL, 
    compress = TRUE, #method to compressing the file
    nb.cpu = 16,
    build.clamping.mask = TRUE,
    seed.val = 1
  )
}

# myBiomodEF
# plot(myBiomodEF@models.projected[2])


# Projecting ensemble models on 1km Europe rasters ---------------------------
run_code <- TRUE
if(run_code){
  
  setwd("Pine Marten/Biomod")
  
  myBiomodEF <- BIOMOD_EnsembleForecasting(
    bm.em = myBiomodEM,
    bm.proj = myBiomodProj_EU,
    proj.name = 'em_eu_lb',
    #new.env = myExpl, #these are the 1km europe rasters
    #new.env.xy = NULL,
    models.chosen = 'all', 
    metric.binary = 'TSS', #vector specifying the names of evaluation metrics and associated thresholds to transform the probabilities of presence into presence and absence. Probably TSS. 
    metric.filter = NULL, 
    compress = TRUE, #metod to compressing the file
    nb.cpu = 16,
    build.clamping.mask = TRUE,
    seed.val = 1
  )
  
}


# Species response curves ---------------------------------
?bm_PlotResponseCurves
response_curves <- bm_PlotResponseCurves(bm.out = myBiomodEM)
land_cover_names <- read.csv("Pine Marten/Environmental rasters/land_cover_names.csv", header=T)

png(filename = "Pine Marten/Other outputs/response_curves.png", width=2000, height=1800, units = "px", res=300)

p1 <- ggplot(subset(response_curves$tab, 
                    expl.name != "land_cover" & 
                      pred.name == "Martes1_EMmeanByTSS_mergedData_mergedRun_mergedAlgo"), aes(x = expl.val, y = pred.val)) + 
  geom_line(col = "black") + 
  labs(title = NULL, x = NULL, y = "Suitability") + 
  facet_wrap(~expl.name, scales = "free_x", labeller = labeller(expl.name = c("elevation" = "Elevation", 
                                                                              "canopy_height" = "Canopy height",
                                                                              "forest_density" = "Forest density",
                                                                              "ndvi" = "NDVI",
                                                                              "slope" =  "Slope",
                                                                              "tpi" = "TPI",
                                                                              "land_cover" = "Land cover"))) + 
  theme_minimal() + 
  theme(legend.position = "none",
        axis.title.y = element_text(hjust=-0.05),
        panel.border = element_rect(colour = "black", fill = NA, linewidth = 1))


p2 <- ggplot(subset(response_curves$tab, expl.name == "land_cover" & 
                      pred.name == "Martes1_EMmeanByTSS_mergedData_mergedRun_mergedAlgo"), aes(x = as.factor(expl.val), y = pred.val)) + 
  geom_col(fill = "lightgrey") + 
  facet_wrap(~expl.name, labeller = labeller(expl.name = c("land_cover" = "Land cover"))) + 
  labs(title = NULL, x = "Variable values", y = "") + 
  scale_x_discrete(labels=land_cover_names$land_cover_name) + 
  #ylim(limits = c(0,layer_scales(p1)$y$range$range[2])) + 
  theme_minimal() + 
  theme(legend.position = "none",
        panel.border = element_rect(colour = "black", fill = NA, linewidth = 1),
        axis.text.x = element_text(angle = 45, vjust = 1, hjust=1))

ggarrange(p1,p2,ncol=1,nrow=2, heights=c(2,1.9))

dev.off()
